# Valid and powerful group statistics for decoding accuracy: Information Prevalence Inference using the i-th order statistic (*i*-test)

**DOI:** 10.1101/578930

**Authors:** Satoshi Hirose

## Abstract

In fMRI decoding studies using pattern classification, a “second-level” group statistical test is typically performed after “first-level” decoding analyses for individual participants. Neuroscientists often test the mean decoding accuracy across participants against the chance-level accuracy (e.g., one-sample Student *t*-test) to verify whether brain activation includes information about the label (i.e., cognitive content). However, Allefeld et al., (2016) notified that positive results for such tests only indicate that “there are some people in the population whose fMRI data carry information about the experimental condition.” Thus, such tests cannot provide inferences about the trend of the majority. They proposed an alternative method, in which prevalence inference is implemented. In this study, I extended their method and propose statistical test “information prevalence inference using the i-th order statistic (*i*-test)”. Compared with the method proposed in the previous study, *i*-test has high statistical power to provide an inference regarding major population trends. In *i*-test, the i-th lowest sample decoding accuracy (the i-th order statistic) is compared to the null distribution to test whether the proportion of the higher-than-chance decoding accuracy in the population (information prevalence) is higher than the threshold. Thus, a significant result of the *i*-test can be interpreted as “*the majority of the population has information about the label in the brain*.” Numerical calculation identified its high statistical power. Also, theoretical detail is provided, and the use of this method in an fMRI decoding study is demonstrated.

Allefeld, C., Görgen, K., Haynes, J.D., 2016. Valid population inference for information-based imaging: From the second-level t-test to prevalence inference. Neuroimage.

## 1. Introduction

An increasing number of fMRI studies use multi-voxel pattern classification analysis, where the cognitive content (= label), such as visual input categories (Spiridon and Kanwisher 2002, Haxby et al., 2014, Nishida and Nishimoto 2018), types of movement (Gallivan et al., 2011, Nambu et al., 2015), and the effector of movement (Hirose et al., 2015, Hirose et al., 2018) is predicted (decoded) from fMRI signal patterns. The prediction’s accuracy (decoding accuracy; D-Acc) is regarded as an index of the label information in the brain. In fMRI decoding studies, the D-Acc obtained in the experiment is often compared to the chance-level accuracy, that is obtained when the brain contains no information about the label (e.g., 50% for binary classifications). When the D-Acc is higher than the chance level, the brain, or a particular brain region, contains information regarding cognitive content. In real experiments, the D-Acc for multiple participants is combined to make an inference with respect to the D-Acc’s population distribution.

This study concerns a statistical test for inferring the D-Acc’s population from the experimental results of multiple participants. In analogy with conventional univariate analysis for voxel activity (Friston et al., 1995), the mean D-Acc across participants has been often compared to the chance level by using Student’s *t*-test (e.g. Gilbert and Fung, 2018, Gallivan et al., 2011) or permutation test (Gallivan et al., 2013, Nambu et al., 2015, Stelzer et al., 2013). The principle of these analyses is that the D-Acc sample mean is a good estimator of its population mean, and if the population mean is higher than chance, the brain activation is expected to typically contain cognitive content information. However, this is a misleading interpretation. Indeed, in theory, the expectation of each participant’s D-Acc has to be equal to or higher than the random classifier and cannot be lower than the chance level (Allefeld et al., 2016). Thus, very small proportion of the population with a higher-than-chance D-Acc is sufficient to make a population mean higher than chance. Therefore, the significant results of these tests cannot provide inferences regarding the population trend; they can only infer that “*there are some people in the population whose fMRI data carry information about the experimental condition*” (Allefeld et al., 2016), indicating that such tests fail to infer the population majority trend.

The above-mentioned problem is inevitable for statistical tests of the population mean. One solution is to assess the population information prevalence (i.e., the proportion of the population with a higher D-Acc than chance level). Accordingly, Allefeld et al. proposed a statistical test on the D-Acc, where the population information prevalence is targeted instead of the population mean. This was inspired by other methods evaluating the proportion of the population for second-level (multi-subject) statistical test in fMRI data analyses, such as the dynamic causal model (Stephan et al., 2009), the second-level random effect of univariate analysis (Rosenblatt et al., 2014), and conjunction analysis (Friston et al., 1999). In this method, the null hypothesis is that the “*proportion of the population that has a higher-than-chance D-Acc is not larger than the predetermined prevalence threshold (e.g.*, 0.5).” Thus, if the null hypothesis is rejected, it can be concluded that “*proportion larger than the prevalence threshold (e.g., more than half*) *of the population has label information in the brain activity*”.

The method proposed in Allefeld et al. (2016) theoretically solved the above concern and achieved a meaningful inference regarding the population trend. However, the method has a drawback. Because the lowest D-Acc among the participants is used as the representative value, the method can often fail to capture the correct population characteristics, which leads poor statistical power. This becomes serious particularly for experiments with many participants, because large number of participants increases probability of appearance of a lower outlier D-Acc, which rarely appears for each participant. This leads a counter-intuitive drawback that the statistical power decreases when the number of participants increases (see Section 3.1).

To address this issue, in the present study, I propose the extension of the previous method. The idea is to generalize Allefeld’s method to the *i*-th order statistic (*i*-th lowest D-Acc). Namely, the 2nd order statistic (2nd lowest D-Acc) or higher-order statistic is used as the representative value. As a result, *i*-test becomes tolerant to lower outliers, because the *i*-th order statistic is immune to the existence of *i* − 1 lower outliers. See Section 2.2.3 for the relationship between the methods proposed by Allefeld et al. (2016) and that proposed in this paper. I name the proposed method as “information prevalence inference using the *i*-th order statistic” or *i*-test.

The remainder of the paper is organized as follows: In the next section, the proposed statistical test is explained (Section 2). Then, the advantage of the method is demonstrated by providing numerical results (Section 3). Next, the application is demonstrated using a real fMRI dataset (Section 4). Finally, the results are summarized, and the key characteristics of the method are briefly discussed (Section 5).

## 2 Information prevalence inference using the i-th order statistic (i-test)

In this section, the statistical model is clarified, the problem is defined, and the underlying theory is outlined (Section 2.0). Next, the theoretical detail of *i*-test is explained (Section 2.1). Then, essential theoretical characteristics of *i*-test are introduced, as well as the relationship with the previous method (Section 2.2). Finally, the practical procedure for performing *i*-test is explained (Section 2.3). Section 2.3 is self-contained, to some extent, and contains minimum equations, so that readers can skip 2.1 and 2.2 to see the practical procedure before the theoretical detail.

### 2.0 notations, problem definition, and outline of the calculation

Figure 1A shows the statistical model of the population (Ω) and how experimental results (sample D-Acc in a real experiment) are obtained. Population Ω can be partitioned into two subgroups, Ω_+_ and Ω_−_. People in Ω_+_ have label information in their brain activity and, thus, expectation of D-Acc higher than chance, while people in Ω_−_ do not. When a person with index *n* is randomly chosen from the population, he/she belongs to Ω_+_ (*n* ∈ Ω_+_), with a probability *γ*, or otherwise belongs to Ω_−_ (*n* ∈ Ω_−_). The experiment with *N* participants is expressed as a two-step random sampling. First, *N* participants are randomly sampled from the population independently of each other (“Random sampling of participant”). Each sampled participant is associated with the probability distribution (probability mass function) of the D-Acc (*p*(*a*_*n*_)). Note that in this paper, lowercase *p* is used for probability mass functions, while upper case *P* is used for probabilities.

**Figure 1:**
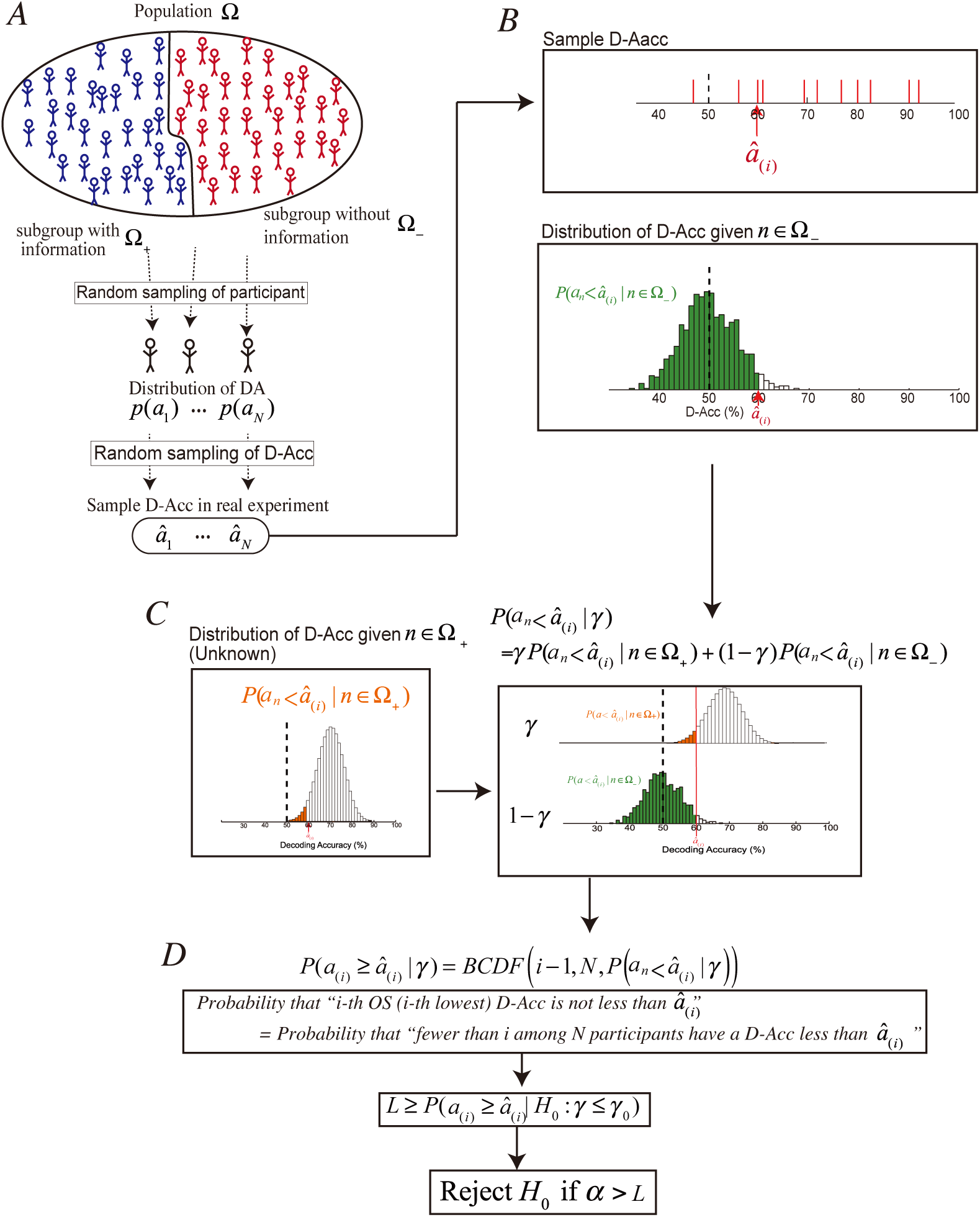
A statistical model and schematic design of *i*-test. **A:** Statistical model. The population (Ω) is composed of two subgroups, Ω_+_, with label information, and Ω_−_, without label information. An experiment is expressed as two-step random sampling; first, each participant is randomly chosen from the population (Random sampling of participant), and then sample D-Acc in the experiment (*â*_*n*_) is randomly chosen from the distribution for each chosen participant (Random sampling of D-Acc). **B:** Values obtained from experimental results. From the experimental results, *i*-th order statistic (*i*-th lowest D-Acc; *â*_(*i*)_ is chosen (top). Also, we estimate the probability distribution given that a participant does not have label information (*p*(*a*_*n*_|*n* ∈ Ω_−_); bottom). Green bars indicate *P*(*a*_*n*_ < *â*_(*i*)_ |*n* ∈ Ω_−_). **C:** Left: The probability distribution given that a participant has label information (*p*(*a*_*n*_|*n* ∈ Ω_+_)). Orange bars indicate *P*(*a*_*n*_ < *â*_(*i*)_ |*n* ∈ Ω_+_), whose value is unknown. Right: the probability that a participant provides D-Acc less than *â*_*n*_ given *γ*, i.e. *P*(*a*_*n*_ < *â*_(*i*)_ |*γ*), can formulate as the weighted sum of *P*(*a*_*n*_ < *â*_(*i*)_ |*n* ∈ Ω_+_) and *P*(*a*_*n*_ < *â*_(*i*)_ |*n* ∈ Ω_−_) with the weight of the probability that a participant belongs to Ω_+_ (*γ*) and to Ω_−_ (1 − *γ*), respectively. **D**: The probability that “*i-th order statistic is not less than observed one* (*â*_(*i*)_” is equivalent to the probability that “*fewer than i among N participants have a D-Acc less than â*_(*i*)_,” which can be formulated with binomial cumulative distribution function. Then, we can calculate the upper bound (*L*) of the probability that “*i-th order statistic is not less than the observed one, given the null hypothesis*:” *P*(*a*_(*i*)_ ≥ *â*_(*i*)_|H_0_: *γ* ≤ *γ*_0_). If this is smaller than the statistical threshold *α*, then, we reject the null hypothesis and conclude that the population prevalence *γ* is larger than *γ*_0_.

Second, the sample D-Acc (*â*_*n*_) is randomly sampled from the distribution for each participant (Random sampling of D-Acc). The objective of *i*-test is to test whether *γ* is larger than the predetermined threshold *γ*_0_ with the significance threshold *α* from *N* participants’ experimental results.

For the statistical test, we first choose the *i*-th order statistic (*i*-th lowest D-Acc; *â*_(*i*)_ from the sample D-Acc observed in the experiment (Figure 1B top). In addition, we estimate the probability mass function of D-Acc given that the participant does not have label information (*p*(*a*_*n*_|*n* ∈ Ω_−_); Figure 1B bottom). This is achieved by empirical procedure, such as permutation test, or with assumption of distribution of D-Acc, such as binomial distribution (See Section 2.2.2). Without estimating the D-Acc probability distribution given a person has label information (*p*(*a*_*n*_|*n* ∈ Ω_+_); Figure 1C left), we formulate the probability that “*a participant provided a D-Acc less than â*_(*i*)_” under a given *γ* (*P*(*a*_*n*_ < *â*_(*i*)_|*γ*); Figure 1C right) as a weighted sum of *P*(*a*_*n*_ < *â*_(*i*)_ |*n* ∈ Ω_+_) and *P*(*a*_*n*_ < *â*_(*i*)_ |*n* ∈ Ω_−_) with the weight of the probability that a participant belongs to Ω_+_ (*γ*) and to Ω_−_ (1 − *γ*), respectively. Then, the probability that “*i-th order statistic is not less than observed one* (*â*_(*i*)_” under a given *γ* is formulated (*P*(*a*_(*i*)_ ≥ *â*_(*i*)_|*γ*); Figure 1D top). This is equal to the probability that “*fewer than i among N participants have a D-Acc less than â*_(*i*)_.” The equivalence of these two events is easily realized by finding that *i*-th lowest D-Acc is not less than *â*_(*i*)_, if and only if the number of participants with D-Acc less than *â*_(*i*)_ is fewer than *i*. Although we cannot calculate the probability directly, we can calculate its upper bound (*L*) under the null hypothesis of “*γ is at or less than the predetermined threshold γ*_0_” (H_0_: *γ* ≤ *γ*_0_). Then, when the upper bound is smaller than *α*, we can reject the null hypothesis and prove that *γ* is larger than the predetermined threshold *γ*_0_ with the significance threshold *α*.

The notations are tabulated in Table 1.

**Table 1:** Notations used in the present study.

|  |
| --- |
| Predetermined Constant |
| $N$: Number of participants |
| $\gamma_0$: prevalence threshold |
| $\alpha$: statistical threshold |
| $i$: rank of order statistic |
| Unknown population value |
| $\gamma$: population information prevalence, i.e. $P(n \in \Omega_+)$ |
| Population |
| $\Omega$: whole population |
| $\Omega_+$: population subgroup with label information |
| $\Omega_-$: population subgroup without label information |
| Probabilistic Variable |
| $a_n$: D-Acc of a participant with index $n$ |
| $a_{(i)}$: $i$-th order statistic of D-Acc from $N$ participants |
| Experimental results |
| $\hat{a}_n$: observed D-Acc of participants in the experiment ($n = 1 \dots N$) |
| $\hat{a}_{(i)}$: the $i$-th order statistic of observed D-Acc in the experiment |
| Function |
| $BPDF(k, K, \rho) = {}_K C_k \rho^k (1 - \rho)^{(K-k)}$ : binomial probability mass function |
| $BCDF(k, K, \rho) = \sum_{h=0}^k BPDF(h, K, \rho)$: binomial cumulative probability mass function |

### 2.1 Theoretical details

As noted above, the objective of *i*-test is to test whether *γ* is larger than the predetermined threshold *γ*_0_. Thus, the alternative hypothesis of H_1_: *γ* > *γ*_0_ is set, and the null hypothesis is its negation H_1_: *γ* ≤ *γ*_0_. Under this null hypothesis, we evaluate whether the *i*-th lowest sample D-Acc *â*_(*i*)_ is so high that it rarely (with a probability less than the significance threshold *α*) occurs.

This calculation is conducted in four steps. First, we formulated the probability that “*each participant’s D-Acc is less than â*_(*i*)_” under a given *γ* (*P*(*a*_*n*_ < *â*_(*i*)_|*γ*:); eq. (2.2)]. Second, the probability that “*the i-th lowest D-Acc is not less than the one actually observed*” under a given *γ* (*P*(*a*_(*i*)_ ≥ *â*_(*i*)_|*γ*:) is formulated (eq. (2.3)). Third, the lower bound (*Q*_*n*_) of the probability that “*each participant’s D-Acc is less than â*_(*i*)_” under the null hypothesis (*P*(*a*_*n*_ < *â*_(*i*)_|H_0_: *γ* ≤ *γ*_0_:) is calculated (eqs. (2.6), (2.7)). Fourth, the upper bound (*L*) of the probability that “*the i-th lowest D-Acc is not less than the one actually observed*” under the null hypothesis (*P*(*a*_(*i*)_ ≥ *â*_(*i*)_|H_0_: *γ* ≤ *γ*_0_:) is calculated (eqs. (2.9), (2.10)). And finally, *L* is compared to *α*.

#### Formulation of *P*(*a*_*n*_ < *â*_*(i)*_|*γ*)

Considering that a participant belongs to Ω_+_ with the probability *γ, p*(*a*_*n*_|*γ*) is formulated as follows:

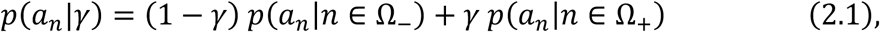

where the D-Acc probability distribution (probability mass function) given a person has and doesn’t have label information is expressed as *p*(*a*_*n*_ |*n* ∈ Ω_+_) and *p*(*a*_*n*_ |*n* ∈ Ω_−_), respectively. From Eq. (2.1), the probability that “*D-Acc of a participant is less than â*_(*i*)_” is formulated as follows:

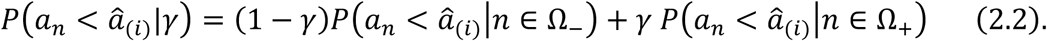

Here, we assume the identical distribution among participants, i.e.

*P*(*a*_*n*_ < *â*_(*i*)_ |*n* ∈ Ω_+_) and *P*(*a*_*n*_ < *â*_(*i*)_ |*n* ∈ Ω_−_) are independent of *n*, and therefore, *P*(*a*_*n*_ < *â*_(*i*)_|*γ*) is independent of *n* (see Appendix A for the relaxation of this assumption).

#### Formulation of *P*(*a*_*(i)*_ ≥ *â*_*(i)*_|*γ*)

Then, we formulate the probability that the “*i-th lowest D-Acc is not less than â*_(*i*)_” under a given *γ* as follows;

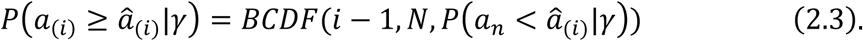

This is because the probability that “*i-th lowest D-Acc is not less than â*_(*i*)_” is equivalent to the probability that “fewer than *i among N participants have a D-Acc less than â*_(*i*)_.”

#### Lower bound of *P*(*a*_i_ < *â*_*(i)*_|H_0_: *γ* ≤ *γ*_0_)

By only noting the basic nature of probability, i.e. *P*(*a*_*n*_ < *â*_(*i*)_|*n* ∈ Ω_+_) ≥ 0, we easily find the lower bound of *P*(*a*_*n*_ < *â*_(*i*)_|*γ*) from eq. (2.2) as

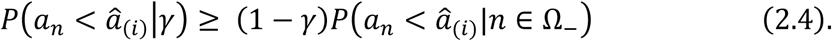

Furthermore, under the null hypothesis H_0_: *γ* ≤ *γ*_0_, we can find the lower bound of the probability as

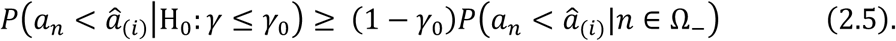

This is because the right side of eq. (2.4) is nondecreasing function of *γ*. Here, we define *Q*_*n*_ as the right side of eq. (2.5), i.e.

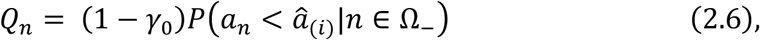

and we can reformulate eq. (2.5) as

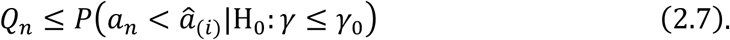

#### Upper bound of *P*(*a*_*(i)*_ ≥ *â*_*(i)*_|H_0_: *γ* ≤ *γ*_0_)

Because *BCDF*(*i* − 1, *N, P*(*a*_*n*_ < *â*_(*i*)_|*γ*)) in eq. (2.3) monotonically decreases with *P*(*a*_*n*_ < *â*_(*i*)_|*γ*), and *P*(*a*_*n*_ < *â*_(*i*)_|H_0_) *γ* ≤ *γ*_0_: is bounded below with *Q*_*n*_ [Eq. (2.7)], we find that *P*(*a*_(*i*)_ ≥ *â*_(*i*)_|H_0_) *γ* ≤ *γ*_0_: is bounded above as follows;

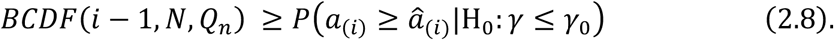

Here, we define *L* as the left side of eq. (2.8), i.e.

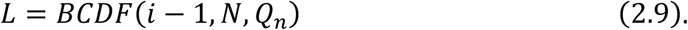

and we can reformulate eq. (2.8) as

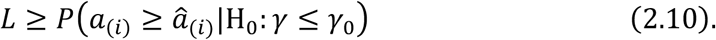

If we know *P*(*a*_*n*_ < *â*_(*i*)_|*n* ∈ Ω_−_), we can calculate *Q*_*n*_ with Eq. (2.6), and consequently, we can calculate *L* with Eq. (2.9). When *L* is smaller than the statistical threshold *α*, it is proven that *P*(*a*_(*i*)_ ≥ *â*_(*i*)_H_0_: *γ* ≤ *γ*_0_) is smaller than *α* from Eq. (2.10). Thus, we can conclude that the event that the “*i-th lowest sample D-Acc is not less than the observed one â*_(*i*)_” rarely (with the probability of less than *α*) occurs under the null hypothesis. Therefore, we can reject the null hypothesis and accept H_1_: *γ* > *γ*_0_, i.e., “*proportion larger than γ*_0_ *of the population has label information in the brain activity.*”

### 2.2 Theoretical characteristics

#### 2.2.1 Predetermined parameters

*i*-test requires three predetermined parameters: *α* (threshold for statistical significance), *γ*_0_ (threshold for the information prevalence), and *i* (rank of the order statistic). Together with the number of participants *N*, they should satisfy the following inequality;

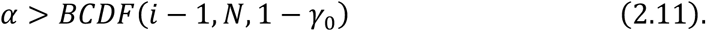

This constraint is derived as follows: From Eq. (2.6) and noting that *P*(*a*_*n*_ < *â*_(*i*)_|*n* ∈ Ω_−_) ≤ 1, *Q*_*n*_ is bounded above with 1 – *γ*_0_. Thus, from Eq. (2.9), and because *L* decreases with *Q*_*n*_, *L* is bounded below as follows:

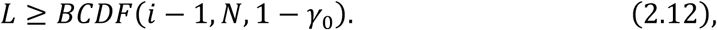

Thus, Eq. (2.11) should be satisfied; otherwise, *L* is always at or larger than *α* and the statistical test never provides a positive result.

See Section 2.3 Step 1 for the empirical guidelines for choosing particular parameter values within this constraint.

#### 2.2.2 Estimation of *P*(*a*_*n*_ < *â*_*(i)*_ |*n* ∈ Ω_−_)

For the calculation of *L*, we need to estimate the probability that “*a participant provided a D-Acc less than â*_(*i*)_” given that a participant does not have label information in the brain, i.e. *P*(*a*_*n*_ < *â*_(*i*)_ |*n* ∈ Ω_−_). This is achieved by estimation of probability mass function *p*(*a*_*n*_|*n* ∈ Ω_−_), by means of empirical procedures such as permutation test (Good, 2000, Ojala and Garriga 2010), or with assumption of its distribution, e.g. binomial distribution. *i*-test can be combined with any kinds of the estimation procedure and this study does not cover the validation of the estimation. But, for the empirical use, I introduce two commonly-used methods.

##### Permutation test

In the permutation test, the labels for the trials are randomly permuted and generate permuted dataset. Then, the decoding analysis is applied to this permuted dataset and calculate the decoding accuracy with the same procedure used in the main analysis. This permutation is repeated multiple times and the distribution of D-Acc from the permutations are regarded as the D-Acc given that there is no label information in the brain.

When *M* permutations are performed for each participant, we calculate D-Acc with each of the permuted data. Then, we estimate *P*(*a*_*n*_ < *â*_(*i*)_ |*n* ∈ Ω_−_) by calculating the proportion of permutations that provide performance less than *â*_(*i*)_,

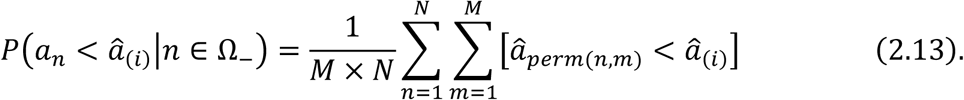

where *â*_*perm*(*n,m*)_ is the D-Acc obtained from the m-th permutation of the n-th participant and Iverson bracket [·] has the value 1 for a true condition and the value 0 for a false condition.

##### Assumption of the binomial distribution

If we assume that the decoder predicted the label randomly and independently in every trials, the D-Acc given the brain activation does not include label information follows the binomial distribution

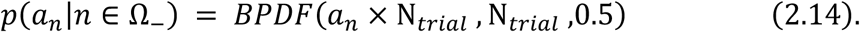

Under this assumption, we estimate *P*(*a* < *â*_(*i*)_ |*n* ∈ Ω_−_) as

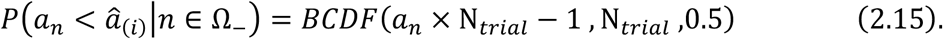

#### 2.2.3 Relation with the previous method in Allefeld et al., (2016)

By fixing *i* = 1, Eq. (2.9) becomes simply the *N*-th power of 1 – *Q*_*n*_;

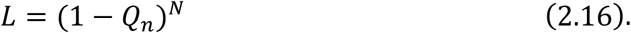

By doing so, *i*-test comes down to the method previously proposed by Allefeld et al., (2016), referred to as the “Permutation-based information prevalence inference using the minimum statistic.” Hereafter, I term this as *i*-test-one.

#### 2.2.4 Possible extension for relaxing the identical distribution among participants

I assume the identical distribution of D-Acc among participants for the derivation of *i*-test. But strictly speaking, this assumption is often broken in a real fMRI situation. Therefore, I propose a possible extension of *i*-test with the relaxation of this assumption. See Appendix A for the derivation of this extension.

### 2.3 Practical Procedure

The procedure for performing the *i*-test is constructed with 1) set three predetermined parameters: *α* (statistical threshold), *γ*_0_ (prevalence threshold) and *i* (rank of the order statistic), 2) identify *â*_(*i*)_ and estimate the probability that “*a participant provided a D-Acc less than â*_(*i*)_” given that a participant does not have label information in the brain, i.e. *P*(*a*_*n*_ < *â*_(*i*)_ |*n* ∈ Ω_−_) and 3) calculate the upper bound of the “*i-th lowest sample D-Acc is at or higher than observed one â*_(*i*)_;” *P*(*a*_(*i*)_ ≥ *â*_(*i*)_|H_0_) *γ* ≤ *γ*_0_: and compare it with *α*.

#### Step 1: Determine the values of the three predetermined parameters

The *i*-test’s positive results can test the hypothesis that “*proportion larger than γ*_0_ *of the population has label information in the brain activity*” with a significant threshold of *α*. In the first step of the *i-*test, we determined the three parameters; *α* (the threshold for statistical significance), *γ*_0_ (the threshold for the information prevalence) and *i* (rank of the order statistic).

Together with the number of participants *N*, The combination of parameters is constrained by eq. (2.11), which determines the upper bound of *i* (*i*_*max*_) when the other parameters, *α*, and *γ*_0_ are fixed. Value of *i*_*max*_ can find by means of numerical calculation of eq. (2.11) implemented as `check_imax.m`.

Under this constraint, there is no universal optimal set of parameter values because it depends on unknown population characteristics. Here I provide a guideline for selecting the parameters. This is not a solid recommendation but suggested as a first choice. First, *γ*_0_ = 0.5 may be intuitively acceptable because we can infer that “*a majority (more than half*) *of the population has label information*,” when we achieve significant result and it is, at present, the standard for existing statistical methods on population prevalence (Friston et al., 1999; see also Discussion in Allefeld et al., 2016). In addition, *α* = 0.05 before the correction for multiple comparisons may be the primary choice because it is the most commonly used in many statistical tests published in neuroscience papers.

As for *i*, if the sufficiently large effect size can be expected and the number of trials is sufficiently large, I recommend to use the largest available *i* (See Section 3; *i*_*max*_). This is because larger *i* can provide higher statistical power, because *i*-test is tolerant to the existence of (*i* − 1) lower outliers (Section 3). However, if the expected effect size and/or the number of trials is relatively small, the statistical power does not monotonically increase with *i* (Appendix C). To decide the value in the empirical situation, I provide a MATLAB program `decide_i.m`, which calculates the expected statistical power for each *i* (Supplementary Figure S4).

#### Step 2: Identify â_(i)_ and estimate *P*(*a*_*n*_ < *â*_*(i)*_ |*n* ∈ Ω_−_)

Then, we identify the value of *â*_(*i*)_ from experimental results. Also, we estimate *P*(*a*_*n*_ < *â*_(*i*)_ |*n* ∈ Ω_−_) by empirical test such as permutation test, or assume that it follows a certain distribution, e.g. binomial distribution (Section 2.2.2).

#### Step 3: Calculate *L* and compare it with *α*

Finally, by substituting the estimation of *P*(*a*_*n*_ < *â*_(*i*)_ |*n* ∈ Ω_−_) to Eq. (2.6) and (2.9), we can calculate the upper bound (*L*) of the probability that the “*i-th order statistic is at or higher than the observed one* (*â*_(*i*)_.” If *L* < *α*, we reject the null hypothesis H_0_: *γ* ≤ *γ*_0_ and conclude that *γ* is larger than the predetermined threshold *γ*_0_ with the significance threshold *α*. The calculation of *L* is implemented in `itest.m.`

## 3 Numerical calculation from artificial data

We designed *i*-test to improve tolerance of the lower outliers compared to when we fix *i* = 1 (*i*-test-one) as proposed by Allefeld et al., (2016). “Lower outlier” here is defined as a low D-Acc that appears for each participant with small probability. Intuitively, if one lower outlier exists in the experimental results, the order statistic *â*_(*i*)_ becomes the lower outlier and the statistical test fails to achieve a significant result even when most of the population has brain activity associated with the label (*n* ∈ Ω_+_). The risk of appearance of one lower outlier becomes larger with the increasing number of participants. Thus, we extended the method so that the second and higher-order statistic can be candidates to make *i*-test tolerant to lower outliers. In this section, I provide an example demonstrating this benefit of *i*-test.

Here, I only provide the numerical calculation results of the probability of the significant result (*P*_*sig*_). See Appendix B for detail of the calculation of *P*_*sig*_.

Note that although the above discussion implies that the largest *i* (*i*_*max*_) is always best, this is true only when the effect size and the number of trials are sufficiently large (Appendix C).

### 3.1 Definition of artificial experimental results

Here, we consider artificial data, where each participant performed *N*_*trial*_ trials with a binary choice (chance level = 50%). The participants have label information (*n* ∈ Ω_+_) with a certain probability *γ* and otherwise, do not have label information (*n* ∈ Ω_−_). We assume that the decoder for participants with label information (*n* ∈ Ω_+_) predicted the label correctly with probability of *P*_*correct* +_ (> chance level), while when *n* ∈ Ω_−_, the decoder for the participant predicted the label in each trial randomly. Also, we assume the independency of each trials. Therefore, the D-Acc for *n* ∈ Ω_+_ and *n* ∈ Ω_−_ follows binomial distributions, i.e. *p*(*a*_*n*_|*n* ∈ Ω_+_) = *BPDF*(*a*_*n*_ *× N*_*trial*_, *N*_*trial*_, *P*_*correct*+_) and *p*(*a*_*n*_|*n* ∈ Ω_−_) = *BPDF*(*a*_*n*_ *× N*_*trial*_, *N*_*trial*_, 0.5). Note that strictly speaking, the cross validated decoding accuracies does not perfectly follow binomial distribution in the real fMRI decoding studies, because independency across trials are not fully met (Jamalabadi et al., 2016). In addition, we assume that *P*(*a*_*n*_ < *â*_(*i*)_|*n* ∈ Ω_−_) is correctly estimated as *BCDF*(*â*_(*i*)_ *× N*_*trial*_ − 1, *N*_*trial*_, 0.5).

The threshold for statistical significance is fixed at 0.05 (*α* = 0.05) and the prevalence threshold is fixed at 0.5 (*γ*_0_ = 0.5). The results are generally replicated with different values of the prevalence threshold (*γ*_0_ = 0.1, 0.0 and 0.*7*; Supplementary Figure S1, S2 and S3).

### 3.2 Results of numerical calculation

First, I fixed the parameters as follows *N*_*trial*_ = 1000, *P*_*correct*+_ = 0.9 and *γ* = 0.8, and I increased the number of participants (*N*) from 5 to 50 and evaluated the probability of significant results (Figure 2A). Note that when *N* < *5*, the predetermined parameters cannot satisfy the constraint [Eq. (2.11)]; therefore, we cannot apply *i*-test.

**Figure 2:**
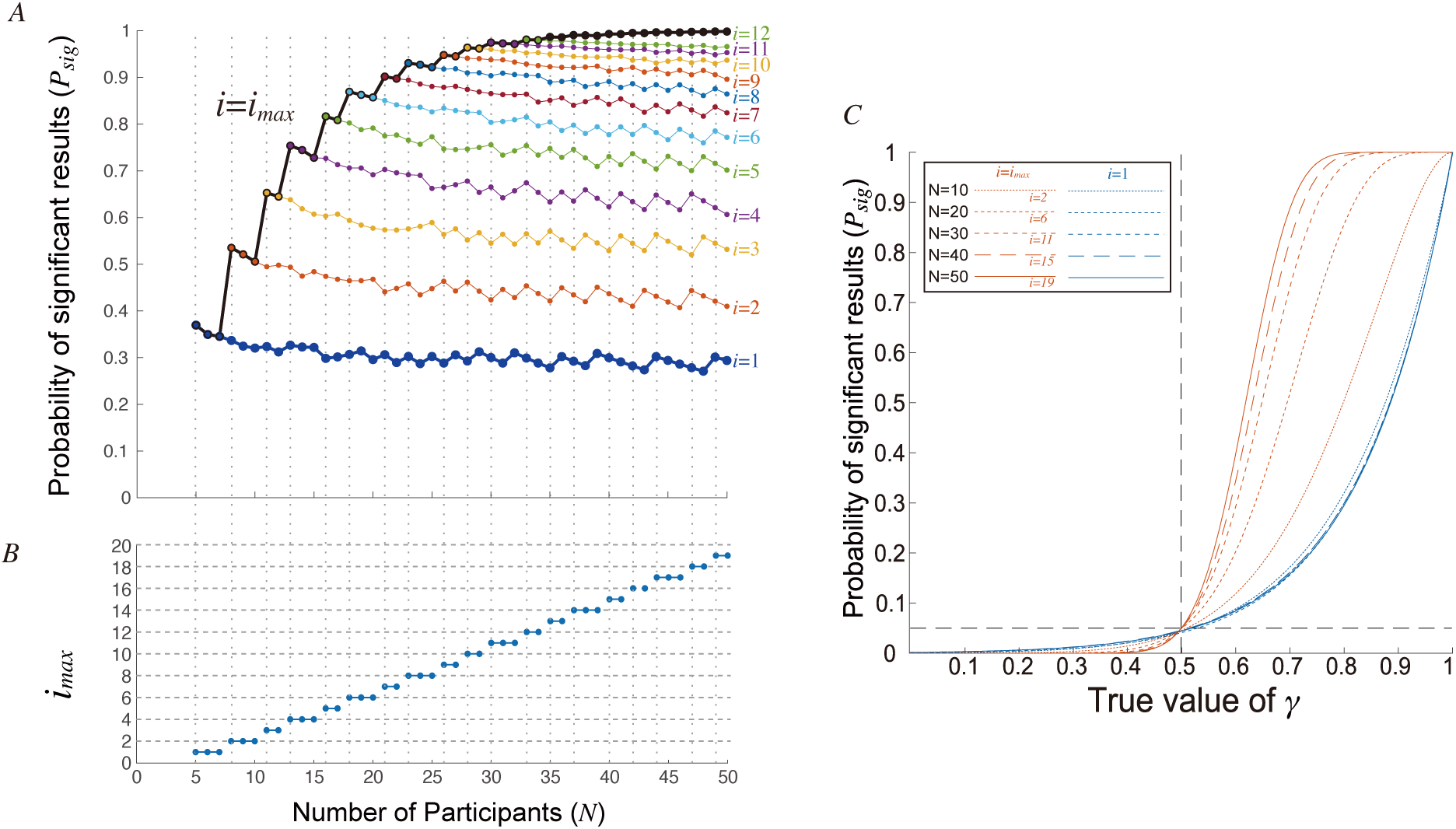
Result from the numerical calculation. The fixed parameters for this analysis are *N*_*trial*_ = 1,000, *P*_*correct*+_ = 0.9, *γ* = 0.8, *α* = 0.05 and *γ*_0_ = 0.5. **A**: The probability of the significant result (*P*_*sig*_) is plotted against the number of participants (*N*). The blue thick line indicates the result of *i*-test-one (*i* = 1). The colored lines indicate the result from fixed *i* for *i* = 1 … 12. The black thick line indicates the result from *i*-test with a maximum available value of *i* (*i*_*max*_). The vertical black dotted lines indicate the point of *i*_*max*_ change. When *N* < *5*, the *i*-test is not available because Eq. (2.9) cannot be satisfied with any *i*. **B**: The value of *i*_*max*_ against *N*. **C**: The probability of a significant result (*P*_*sig*_) are plotted against the value of *γ*. Red lines represent results from *i*-test with a maximum available value of *i* (*i* = *i*_*max*_) and blue lines represent results for *i*-test-one (*i* = 1). The number of participants is 10, 20, 30, 40 or 50 and *i*_*max*_ is 2, 6, 11, 15 or 19, respectively. The vertical and horizontal dotted black lines indicate the prevalence threshold *γ*_0_ = 0.5 and the statistical threshold *α* = 0.05, respectively. When *γ* > 0.5 (right side from the vertical dotted line), the alternative hypothesis H_1_: *γ* > *γ*_0_ is true and thus, *P*_*sig*_ indicates statistical power and is ideally close to 1, while *γ* ≤ 0.5, the null hypothesis H_0_: *γ* ≤ *γ*_0_ is true and thus, *P*_*sig*_ indicates a false alarm ratio and should be smaller than *α*.

The probability of a significant result (*P*_*sig*_) of *i*-test-one (*i* = 1) gradually decreases as the number of participants (*N*) increases from 5 to 50 (thick blue line in Figure 2A). Also, when *i* is fixed at a certain value (colored thin lines in Figure 2A), the probability of a significant result gradually decreases. However, larger *i* provided higher *P*_*sig*_ and the maximum available value of *i* (*i*_*max*_) also increased with *N* (Figure 2B). Thus, by using *i*_*max*_, *i*-test can improve the statistical power with increase of *N* (black thick line in Figure 2A). The reduction of D-Acc of *i*-test-one is because the probability of contamination of lower outlier increases when the number of participants increases. In contrast, the improvement of *i*-test with *i*_*max*_ is because *i*-test is tolerant to existence of *i* − 1 lower outliers (Appendix B).

Then, we evaluated the effect of *γ* on *P*_*sig*_. *γ* was changed from 0 to 1 with 0.01 step (Figure 2C). Note that when *γ* is at or below the prevalence threshold *γ*_0_ (left side of the Figure 2C), the null hypothesis is true and *P*_*sig*_ indicates false positive ratio that should be suppressed below the statistical threshold (*α*). Again, I fixed *N*_*trial*_ = 1000, and *P*_*correct*+_ = 0.9. Figure 2C shows the results from *N* = 10, 20, 00, 40, 50. I found that both *i*-test-one and *i*-test with *i* = *i*_*max*_ could suppress the false alarm ratio below the statistical threshold (Figure 2C left). Also, I found that the statistical power (*P*_*sig*_ at *γ* > *γ*_0_; Figure 2C right) of *i*-test with *i*_*max*_ is almost always higher than the *i*-test-one, while the false alarm ratio (*P*_*sig*_ at *γ* ≤ *γ*_0_; left side of the figure 2C) of *i*-test with *i*_*max*_ is lower for almost all *γ* ≤ *γ*_0_. Exception was observed only when the value of *γ* is close to *γ*_0_ (*γ*_0_– 0.04 < *γ* ≤ *γ*_0_ *+* 0.01). This was replicated with the other values of *N* from 5 to 50 (Results not shown).

## 4 Application to empirical fMRI data

Here, we provide an example of its application to a real fMRI dataset, which is reported in Hirose et al., 2018.

### 4.1 Experiment and analyses procedures

For experimental details, please refer to the original paper (Hirose et al., 2018). Briefly, 12 participants performed a finger-tapping task using either his/her freely-chosen left- or right-hand finger. Each participant underwent 10 sessions, and each session consisted of 15 trials. Each trial started with a 4-sec decision period, where participants chose either their left or right hand to be moved. Then, after a 4-s delay, the participants moved the selected hand finger. Trials were separated by an inter-trial interval (ITI) of 6 s and, in the previous study (Hirose et al., 2018), we carefully evaluated the effect of the previous trial and concluded that there was no carry over of the brain activity for the previous trial. We predicted the chosen hand from brain activity measured just before the movement execution (latter 2 seconds of the delay).

Although each participant underwent 150 trials in total, there were error trials, in which no finger movement occurred, or the participant moved multiple fingers. The number of error trials was 11 for a participant, 5 for another participant, and 1 for other two participants. No error trials were found in the other eight participants. The error trials were excluded from the decoding analysis. Consequently, brain activation for 139 trials for one participant, 145 for another, 149 for other two and 150 for the other eight participants were used for the decoding analysis.

The preprocessed fMRI volume measured before an action initiation is used to predict the chosen hand. The classifier was trained by the iSLR algorithm (Hirose et al., 2015) on voxel activities in the whole brain (31,957 ± 1785). Classification accuracy was evaluated by the leave-one-session-out cross-validation procedure, in which we trained the classifiers using the trials of the nine sessions (training dataset) and used the trials of the remaining session to evaluate the prediction accuracy (test dataset). The balanced accuracy (Brodersen et al., 2010, Brodersen et al., 2012) was calculated from the cross-validation results, because there were bias of the number of trials between labels depending on the bias of the participants’ choice.

Also, I performed the permutation test to estimate *p*(*a*_*n*_|*n* ∈ Ω_−_) with the following procedures. First, the labels (left or right hand) in each session were randomly permuted for each participant. Using this permuted dataset, we then performed the decoding analysis and calculated the decoding accuracy via leave-one-session-out cross validation, using the same procedure as applied to the original (unpermuted) data. To save computational cost, the hyperparameter of the iSLR algorithm (i.e., the number of SLR classifiers) was not re-optimized and the same value as in the analysis without permutation was used in each validation fold. This was repeated 1000 times for each participant, to obtain 1000 decoding accuracies. Then, the decoding accuracies were used to estimate *P*(*a*_*n*_ < *â*_(*i*)_ |*n* ∈ Ω_−_) through eq. (2.13). Note that in each permutation, we randomly shuffled the association between brain activities and labels (left or right hand). Therefore, the number of trials assigned with one label is saved in each session.

The preprocessing, first-level decoding analyses, and permutation test for each participant was performed using multi-voxel pattern classification toolbox (https://github.com/satoshi-hirose/MVPC_toolbox).

Then, I applied *i*-test and *i*-test-one to the experimental results and the permutation test results. For all tests, we set the predetermined thresholds as *α* = 0.05 and *γ*_0_ = 0.5. For the *i*-test, I used *i* = 0, which is the maximum available value that satisfies Eq. (2.11) and was expected to provide highest statistical power when *P*_*correct*+_ > 0.65 (Supplementary Figure 4).

### 4.2 Results

#### *i*-test

The 3rd order statistic was 60.0% (*â*_(3)_ = 60.0+). In the permutation test, the number of permutations with D-Acc lower than *â*_(3)_ was 11,684 (blue bars and green bars in Figure 3) among 12,000 (12 × 1,000) permutations. Therefore, we estimated *P*(*a*_*n*_ < *â*_(*i*)_|*n* ∈ Ω_−_) as 0.97 [11,684/12,000, Eq. (2.13)]. Substituting this value in Eq. (2.6) and Eq. (2.9), we found that *L* = 0.024 and this was smaller than the statistical threshold *α* = 0.05. Thus, the null hypothesis is rejected.

**Figure 3:**
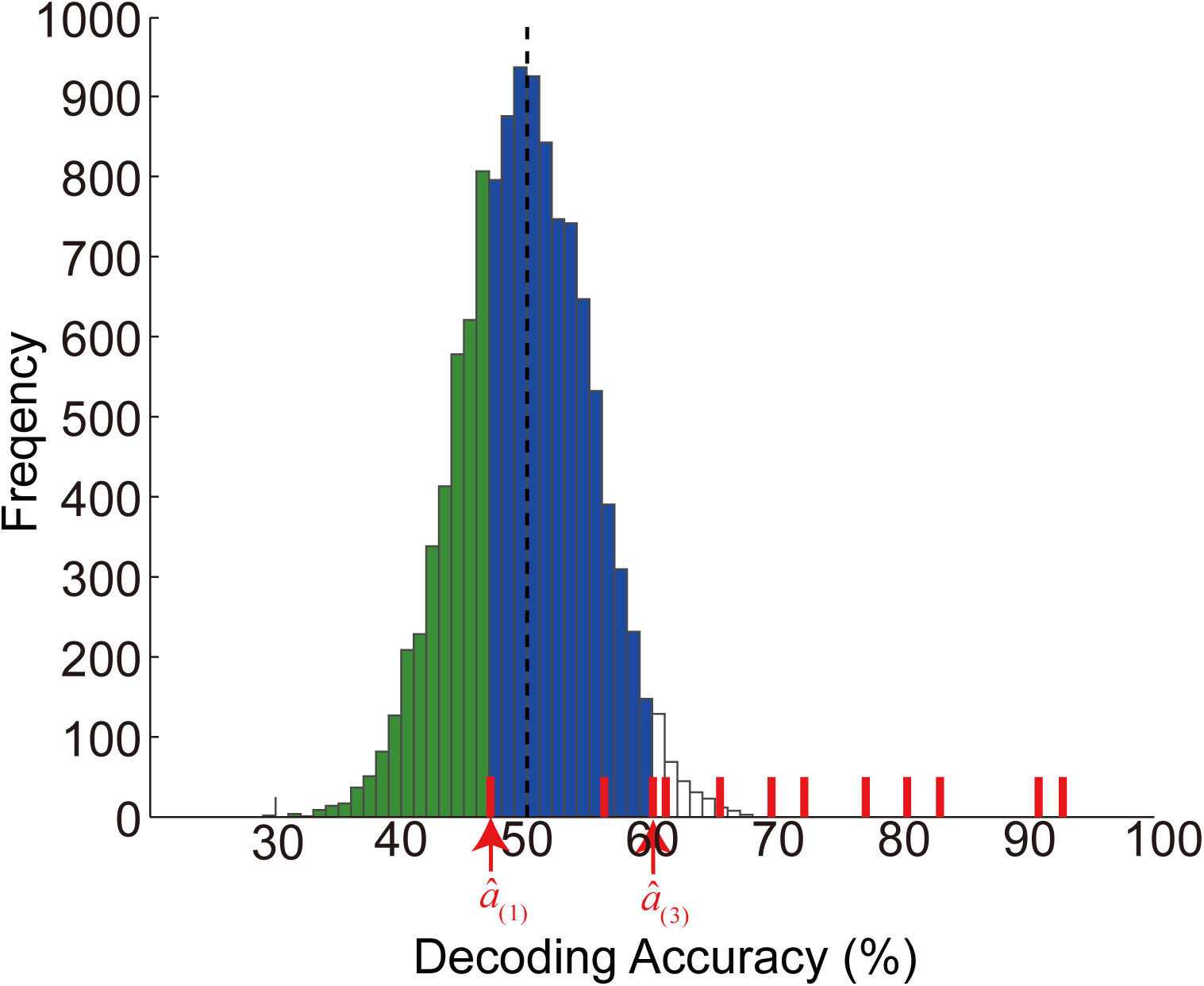
Permutation test results in the empirical experiment for the *i-*test with *i* = 0 and i-test-one (*i* = 1). The histogram indicates the frequency of permuted D-Acc *a*_*perm*(*n,m*)_ from 12,000 permutations (12 participants × 1,000 permutations). Green bars indicate the area of *a*_*perm*(*n,m*)_ < *â*_(1)_, whereas *a*_*defg*(*n,g*)_ < *â*_(3)_ is indicated by the blue bars and green bars.

#### i-test-one

We use the lowest observed D-Acc (*â*_(1)_ = 47.6+) for *i*-test-one. From the permutation results, we found that the D-Acc was lower than 47.6% in 3993 of the 12,000 permutations (green bars in Figure 3). Thus, we estimated *P*(*a* < *â*_(1)_|*n* ∈ Ω_−_) as 0.33. Using Eq. (2.6) and (2.16), we found *L* = 0.11. Thus, *i*-test-one could not reject the null hypothesis.

We cannot know the true population for the empirical data. However, when we look at the distribution of the obtained D-Acc (Red bars in Figure 3), the value of *â*_(1)_ can appears to “lower outlier,” that unfortunately contaminated the results. Thus, we may assume that *i*-test-one failed to achieve significant results because one participant provides low (outlier) D-Acc. And *i*-test could achieve the significant result, because of its tolerance of lower outliers.

## 5. Discussion

In this study, we proposed a novel statistical test for decoding accuracy and named it *i*-test, which is an extension of the method proposed by Allefeld et al. (2016). In contrast to the commonly used statistical test, which compares the population mean D-Acc with the chance level, our method focuses on population information prevalence, i.e., the proportion of the population that has a higher-than-chance D-Acc. The positive result of the statistical test on population information prevalence can infer major population trends as “*the brain activation has label information in proportion larger than γ*_0_ *of the population (e.g., more than half*).” This is in contrast to the fact that the positive result of the statistical test on the population mean D-Acc can only infer that “*there are some people in the population whose fMRI data carry information about the experimental condition*,” because expectation of each participant’s D-Acc should never be lower than the chance level.

A statistical test proposed by Allefeld et al., 2016 (*i*-test-one) also evaluates the population prevalence rather than the population mean. In their method, the minimum statistic is necessarily used for the test. However, as shown in Section 3, using the minimum statistic leads to weakness to lower outliers. This weak point leads to a counter-intuitive drawback: *i*-test-one can reduce its statistical power as the number of participants increases (Figure 2A). This drawback is solved using a higher-order statistic in the *i*-test.

### 5.1 Estimation of *P*(*a*_*n*_ ≥ *â*_*(i)*_ |*n* ∈ Ω_−_)

In the analysis of the real experiment (Section 4, Figure 3), I estimated the distribution of D-Acc given that label information is not included in the brain (*p*(*a*_*n*_|*n* ∈ Ω_−_)), by means of the permutation test, in which labels were permuted within each session before the cross validation. However, there are other candidates for the procedures, e.g. permute label among whole experiment and permute only within training/test session for each cross validation. At present, little is known about the advantage and disadvantage of each procedure.

Also, parametric procedure with assumption of binomial distribution (as I have done in the numerical calculation) can be more appropriate in some situation. For example, when there is limited number of samples, e.g. 4 samples for each label, the maximum number of permutations is 70, which might be insufficient for the empirical estimation of the distribution.

The present study does not cover the validation of these methods. Further study is needed to understand the advantage and disadvantage of the permutation procedures and parametric methods.

## 6. Conclusions

In this study, a novel group-level statistical test for the decoding accuracy named *i*-test is proposed. The advantages of the *i*-test are that the *i*-test 1) can provide a mathematically guaranteed, meaningful population inference; 2) is tolerant to lower outliers and can provide high statistical power.

Although the *i*-test is being introduced as a statistical method for the D-Acc, it is also applicable to other “information-like” measures, such as Mahalanobis distance (Kriegeskorte et al., 2006, Nili et al., 2014), linear discriminant t (Nili et al., 2014) or pattern distinctness D (Allefeld and Haynes, 2014). We expect this study to provide a robust, second-level statistical test for information-based neuroimaging studies.

## 7. Data and code availability

All the analyses in this study can be replicated by MATLAB program codes available at https://github.com/satoshi-hirose/i-test, including implementation of *i*-test and D-Acc obtained from the experiment (Hirose et al., 2018) and from permutation test (1,000 random permutation) for each participant.

## Acknowledgments

I would like to acknowledge Dr. Atsushi Yokoi, Center for Information and Neural Networks (CiNet), National Institute of Information and Communications Technology, and Dr. Isao Nambu, Nagaoka University of Technology, for helpful suggestions on earlier versions of the manuscript. This work was supported by a Grant-in-Aid for Young Scientists (B) (JSPS KAKENHI 16K16649) to the author.

## Abbreviations

D-Acc: decoding accuracy

## Appendix A: Extension for relaxing the assumption of identical distribution among participants

*i*-test can extend to relax the assumption of the identity of the distribution among participants (assumption of i.d.), which is required for most of the existing statistical tests and I used it in *i*-test to derive eq. (2.3). In this extension, we calculate the values of *L* without this assumption.

Strictly speaking, the assumption of i.d. is often broken in a real fMRI situation. For example, in our experiment (Section 4), the number of trials were different among participants, because of existence of error trials and thus, possible values of decoding accuracy *a*_*n*_ should be different. By considering this issue, at least theoretically the relaxation of the assumption should have a merit to provide a good fit to the true population. However, because we have no clear difference in the empirical and numerical analysis and the relaxation has a demerit on computational cost (Supplementary Figure S5), I only present this as a possible extension of *i*-test.

### A1: Derivation

First, I present the derivation of the extended version. Note that eqs. (2.1), (2.2), (2.4), (2.5), (2.6) and (2.7) is derivation for each participant and therefore, we do not need to replace for the extension.

Here, we do not assume the identical distribution of *p*(*a*_*n*_|*n* ∈ Ω_−_) among participants, we cannot express “*i-th lowest D-Acc is not less than â*_(*i*)_” with binomial distribution function as eq.(2.3). Instead, *P*(*a*_(*i*)_ ≥ *â*_(*i*)_|*γ*) is expressed with sum of the probability of the all cases;

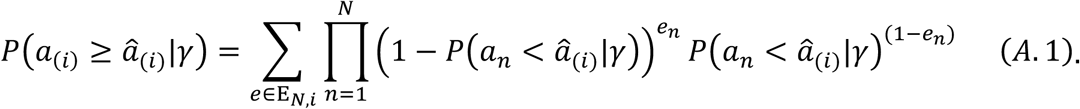

Here, *E*_*N,i*_ is the group of all *N*-dimensional vectors, with elements of 1 and 0, and the number of 0s is less than *i* (the number of 1s is more than *N* − *i*). When a vector *e* is an element of *E*_*N,i*_, *e*_*n*_ is the *n*-th element of *e*. The *N*-dimensional vector *e* represents each combination, and each element of *e* represents whether the D-Acc of the corresponding participant is higher than *â*_(*i*)_ (1) or not (0). For example, when *N* = 0 and *i* = 2, *E*_*N,i*_ = {[1 1 1], [0 1 1], [1 0 1], [1 1 0]} and the first element of *E*_*N,i*_ ([1 1 1]), represents the case that the D-Acc of all three participants are at or higher than *â*_(*i*)_ and [0 1 1] represents the case that the D-Acc of participant #1 is less than *â*_(*i*)_ but the remaining two (#2 and #3) are at or higher than *â*_(*i*)_. The probability of each case is the product of the probability of each participant’s result, e.g., when 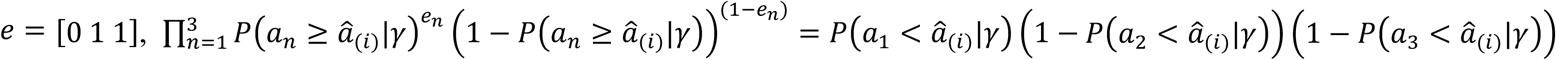.

Then, eqs. (2.8), (2.9) are replaced with

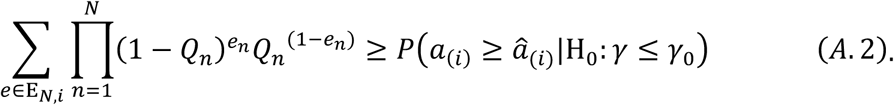

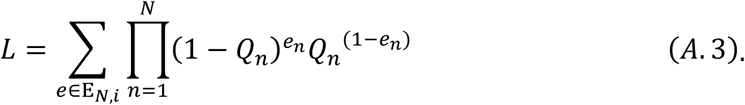

A notice for the derivation of A.2 is that *P*(*a*_(*i*)_ ≥ *â*_(*i*)_|*γ*) in Eq. (A.1) monotonically decreases with each *P*(*a*_*n*_ < *â*_(*i*)_|*γ*). The monotonicity is obvious because when the probability that the “*D-Acc of one of the participants is less than â*_(*i*)_” (*P*(*a*_*n*_ < *â*_(*i*)_|*γ*)) increases, the probability that “*more than N* − *i participants have a D-Acc equal to or higher than â*_(*i*)_” (*P*(*a*_(*i*)_ ≥ *â*_(*i*)_|*γ*)) should decrease. Then, by comparing *L* with *α*, we can perform *i*-test without the assumption of identical distribution among participants.

### A.2 Empirical example

Here, I applied the extended *i*-test to the experimental results used in Section 4 (Hirose et al., 2018). As in Section 4, we set the predetermined parameters as *α* = 0.05, *γ*_0_ = 0.5. and *i* = 0. Unlike the original *i*-test, the estimation of the D-Acc distribution given that a participant does not have label information in the brain (*p*(*a*_*n*_|*n* ∈ Ω_−_)) should be done for each participant. Thus, eq. (2.12) is replaced with the following equation for each participant;

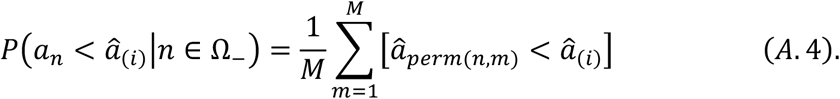

Here, we found that the estimated values ranged from 0.963 to 0.983. By substituting these values into Eqs. (2.6) and (A.3), we found *L* = 0.024, which is similar to the original *i*-test (difference < 0.000002). Thus, this extended version also rejected the null hypothesis.

#### 2.2 Computational cost

A demerit of this extension is its large computational cost. Because elements of *E*_*N,i*_ rapidly increase with *N* and *i*, the calculation of A.3 easily suffers a combination explosion of computational cost. Empirical benchmark test identified that the computation of the extended *i*-test with *i*_*max*_ was completed within 1 s when *N* < 20 and was completed within 1 min when *N* < 28, but it took about 5.5 min for *N* = 00 (Supplementary Figure 6A). Thus, the empirical upper bound of *N* may be about 30. This is contrasted with the original *i*-test that requires computational time that linearly increases with *N* and requires less than 0.03 s for *N* = 1,000 (Supplementary Figure 6B).

## Appendix B: Numerical calculation of *P*_*sig*_

Here I describe the calculation of the probability of the significant result (*P*_*sig*_) reported in Section 3. The outline of the calculation is as follows. First, I introduce the threshold of the *i*-th order statistic; *T*, such that *i*-test provides positive results when and only when *a*_(*i*)_ > *T* (Section B1). The value of T can be derived from the true DA distribution (*γ, p*(*a*_*n*_|*n* ∈ Ω_−_) and *p*(*a*_*n*_|*n* ∈ Ω_+_)) and the predetermined parameters (*α, γ*_0_ and *i*). Note that here, I assume the binomial distribution for *p*(*a*_*n*_|*n* ∈ Ω_−_) and *p*(*a*_*n*_|*n* ∈ Ω_+_). Then, I calculate the probability that *i*-th order statistic is larger than *T* (*P*(*a*_(*i*)_ > *T*); Section B2).

To improve understanding, numerical example with the following parameters are provided in each step; *α* = 0.05, *γ*_0_ = 0.5, *N* = 50, *i*=1, *N*_*trial*_ = 1,000, *P*_*correct*+_ = 0.9 and *γ* = 0.8.

### B1 Introduction of the threshold of *i*-th order statistic; *T*

From the assumptions, *P*(*a*_*n*_ < *â*_(*i*)_|*n* ∈ Ω_−_) is expressed with binomial cumulative distribution function as follows;

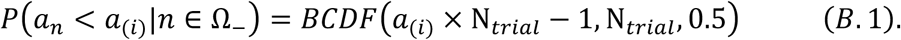

Here, I repeat the equation for *i*-test [i.e., Eq. (2.6) and Eq. (2.9)].

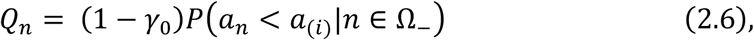

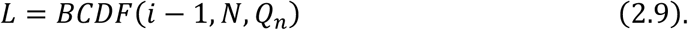

Note that here, the *i*-th order statistic is not a constant but probabilistic variable and therefore, it is expressed without hat. Form eqs. (B.1), (2.6) and (2.9), *L* can be regarded as a (discrete) function of *a*_(*i*)_. *L* decreases with *â*_(*i*)_, because *P*(*a*_*i*_ < *â*_(*i*)_|*n* ∈ Ω_−_) increases with *a*_(*i*)_, *Q*_*n*_ increases with *P*(*a*_*n*_ < *â*_(*i*)_|*n* ∈ Ω_−_) and *L* decreases with *Q*_*n*_. Therefore, we can define the threshold *T*, such that *α* > *L* when and only when *â*_(*i*)_ > *T*. Because the D-Acc is discrete, we can easily perform numeric calculation of *T* from *p*(*a*_*n*_|*n* ∈ Ω_−_), *γ*_0_, *i, N* and *N*_*trial*_, without analytical solution (Supplementary Figure S6).

In the example case with parameters listed above we find *T* = 0.481.

### B2 Calculate the probability that *i*-th order statistic is higher than *T*

By definition, the probability of the significant result is determined as follows:

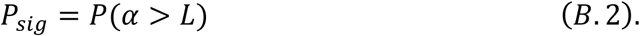

From equivalence of *α* > *L* and *a*_(*i*)_ > *T*, we find

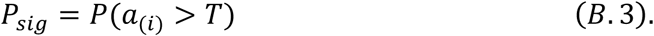

Below, the goal is to calculate *P*(*a*_(*i*)_ > *T*). First, I formulate the probability that “*the D-Acc of a participant is at or less than T*,” as follows;

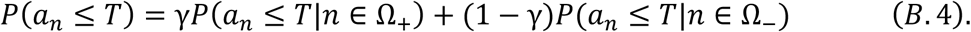

Then, by noting that “*i-th order statistic is higher than the threshold* (*T*)” is identical to “*fewer than i among N participants have a D-Acc at or lower than T*,” *P*(*â*_(*i*)_ > *T*) can be expressed as follows;

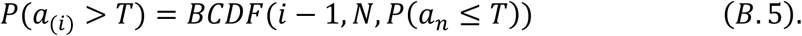

From eq.(B.4) together with true distribution (*p*(*a*_*n*_|*n* ∈ Ω_+_), *p*(*a*_*n*_|*n* ∈ Ω_−_) and *γ*), we can calculate the value of *P*(*a*_*n*_ ≤ *T*). Then, from eq. (B.5), we can calculate *P*(*a*_(*i*)_ > *T*), which is equal to *P*_*sig*_.

In the example, *P*(*a*_*n*_ ≤ *T*|*n* ∈ Ω_+_) and *P*(*a*_*n*_ ≤ *T*|*n* ∈ Ω_−_) was, as follows;

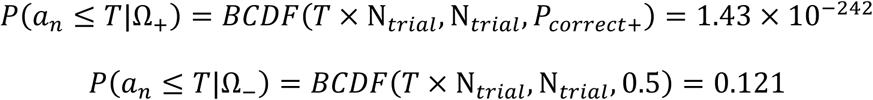

Then, substituting them together with *γ* = 0.8 into eq.(B.4), we find *P*(*a*_*n*_ ≤ *T*) = 0.0242. By substituting it to eq.(B.5) together with *i* = 1 and *N* = 50, we find *P*(*â*_(*i*)_ > *T*: = 0.294.

This means that, although the probability that “*D-Acc of a participant is at or less than T*” *P*(*a*_*n*_ ≤ *T*) is small (3%), the probability that “*D-Acc of <u>no</u> participant among N is less than T*” *P*(*â*_(1)_ > *T*: is only 29%, because there are many (50) participants. As a result, *i*-test-one failed to achieve significant results with probability of more than 70%, although the true *γ* (0.8) is much larger than the prevalence threshold *γ*_0_ = 0.5.

This is solved by using larger *i*. When we use the parameter as *i* = 19 (*i*_*max*_), the threshold T becomes slightly higher (*T* = 0.500) and as a results, the probability that “*the D-Acc of a participant is less than T*,” becomes larger *P*(*a*_*n*_ ≤ *T*) = 0.195. However, with *i* = 19, *i*-test can provide significant result when “*D-Acc of fewer than* 19 *participants among N is less than T.*” This indicates that *i*-test can be tolerant to the 18 “lower outliers” (i.e. participants with D-Acc lower than *T*) and as a result, *i*-test can provide significant result almost 100% (*P*(*â*_(19)_ > *T*) = 0.998). These results can be linked with my design concept of *i*-test for the tolerance of lower outlier. Namely, when we define D-Acc below the threshold *T* as “lower outlier,” the tolerance of lower outlier can be achieved by generalize the *i*-test-one (Allefeld et al., 2016) to *i*-test and as a result, *i*-test can achieve higher statistical power.

## Appendix C: *i*_*max*_ is not always optimal

As demonstrated in the Section 3, larger *i* leads tolerance of lower outlier and consequently can provide higher statistical power. This implies that the largest *i* (*i*_*max*_) is the optimal choice. However, this is not a general fact.

For example, when the probability of the correct decoding for Ω_+_ (*P*_*correct*+_) is low (*P*_*correct*+_ = 0.50), the relation between *i* and *P*_*sig*_ is not monotonical but inverse-U shape (Appendix Figure C.1A). Another example is shown in Figure 3B. When the number of trials is small (*N*_*trial*_ = 10), the relation seems to be more complex; *P*_*sig*_ generally increased with *i* but, decreases when *i* increases from 1 to 2, from 5 to 6, from 10 to 11, from 15 to 16 and from 18 to 19 (Appendix Figure C.1B).

These phenomena can be explained as follows. From the derivation of *P*_*sig*_ (Appendix B), we can find that increase of *i* affect *P*_*sig*_ in two pathways. First (increase effect), when *i* increases, the number of ignorable lower outliers (*a*_*n*_ ≤ *T*) increases (eq.(B.5)). Therefore, it increases the statistical power as demonstrated in Section 3. Second (decrease effect), the threshold *T* can increase as *i* increases and consequently, probability of “lower outlier” *P*(*a*_*n*_ ≤ *T*) can increase. Then, increase of *P*(*a*_*n*_ ≤ *T*) leads decrease of *P*_*sig*_. The decrease effect becomes dominant when *i* is large in the first example with low *P*_*correct*+_. And in the second example, decrease effect appears only at certain points, because of the discreteness of possible values of *T*.

Anyhow, the optimal *i* is decided with the balance between these two effects and the effects depends on the known parameters, such as number of trials, as well as unknown true parameters, such as the probability of the correct decoding for participants belonging to Ω_+_ (*P*_*correct*+_).

However, the parameter *i* should be decided without observation of data to avoid inflation of false positive rate by “cheating effect.” For the purpose, I recommend selecting *i* that maximizes expected statistical power as usually done in “a priori power analysis” (Faul et al., 2009, G*power; http://www.psychologie.hhu.de/arbeitsgruppen/allgemeine-psychologie-und-arbeitspsychologie/gpower.html). For this parameter determination, I provide a program code (`decide_i.m`) that enable us to calculate the statistical power under the assumption of binomial distribution of the decoding accuracy. An example is shown in Supplementary Figure 4.

---

**Figure A1:**
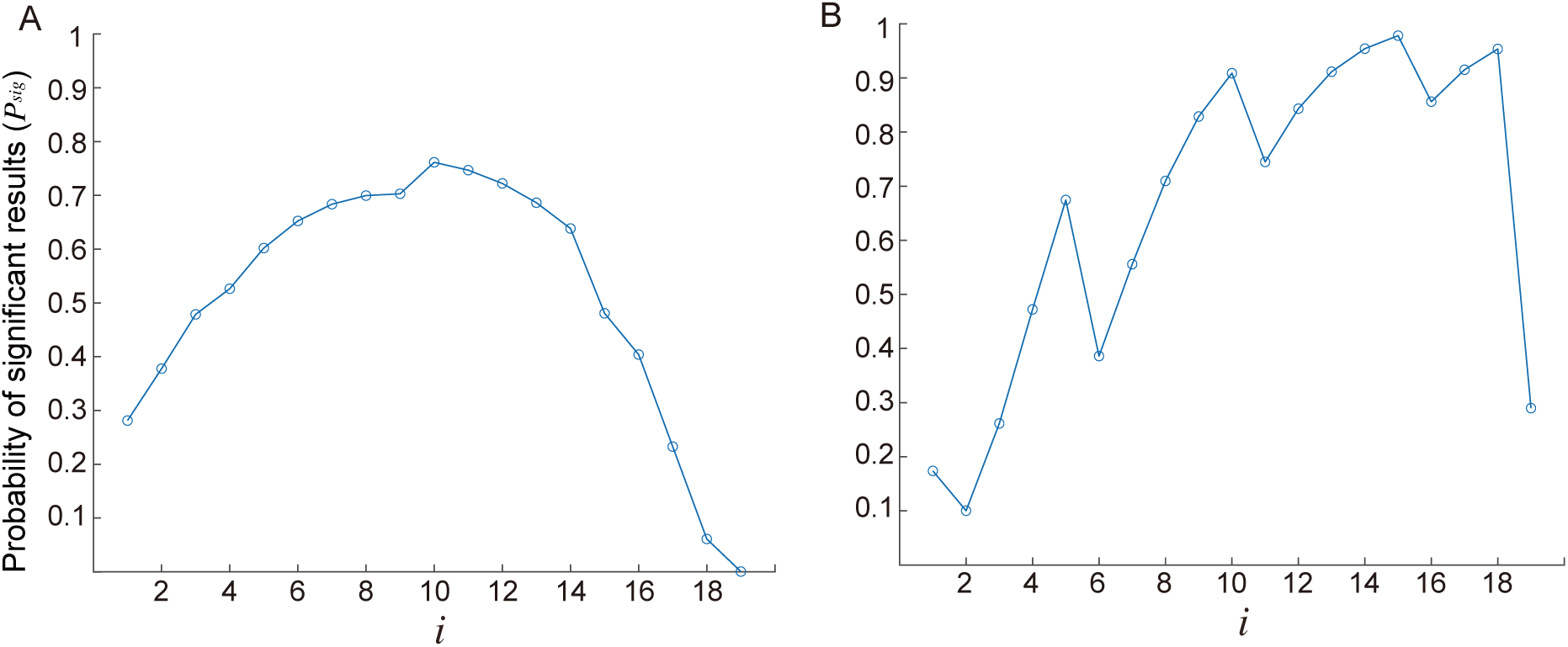
Result from the numerical calculation. **A**: Effect of low effect size (*P*_*correct*+_ = 0.53). The probability of the significant result (*P*_*sig*_) is plotted against the parameter *i*. The number of participants was fixed as *N* = 50. The other fixed parameters were the same as those of Section 3.2 (*N*_*trial*_ = 1,000, *γ* = 0.8, *α* = 0.05 and *γ*_0_ = 0.5). **B**: Effect of small number of trials (*N*_*trial*_ = 10). The probability of the significant result (*P*_*sig*_) is plotted against the parameter *i*. The number of participants was fixed as *N* = 50. The other fixed parameters were same as those of Section 3.2 (*γ* = 0.8, *P*_*correct*+_ = 0.9, *α* = 0.05 and *γ*_0_ = 0.5).

**Supplementary Figure S1:**
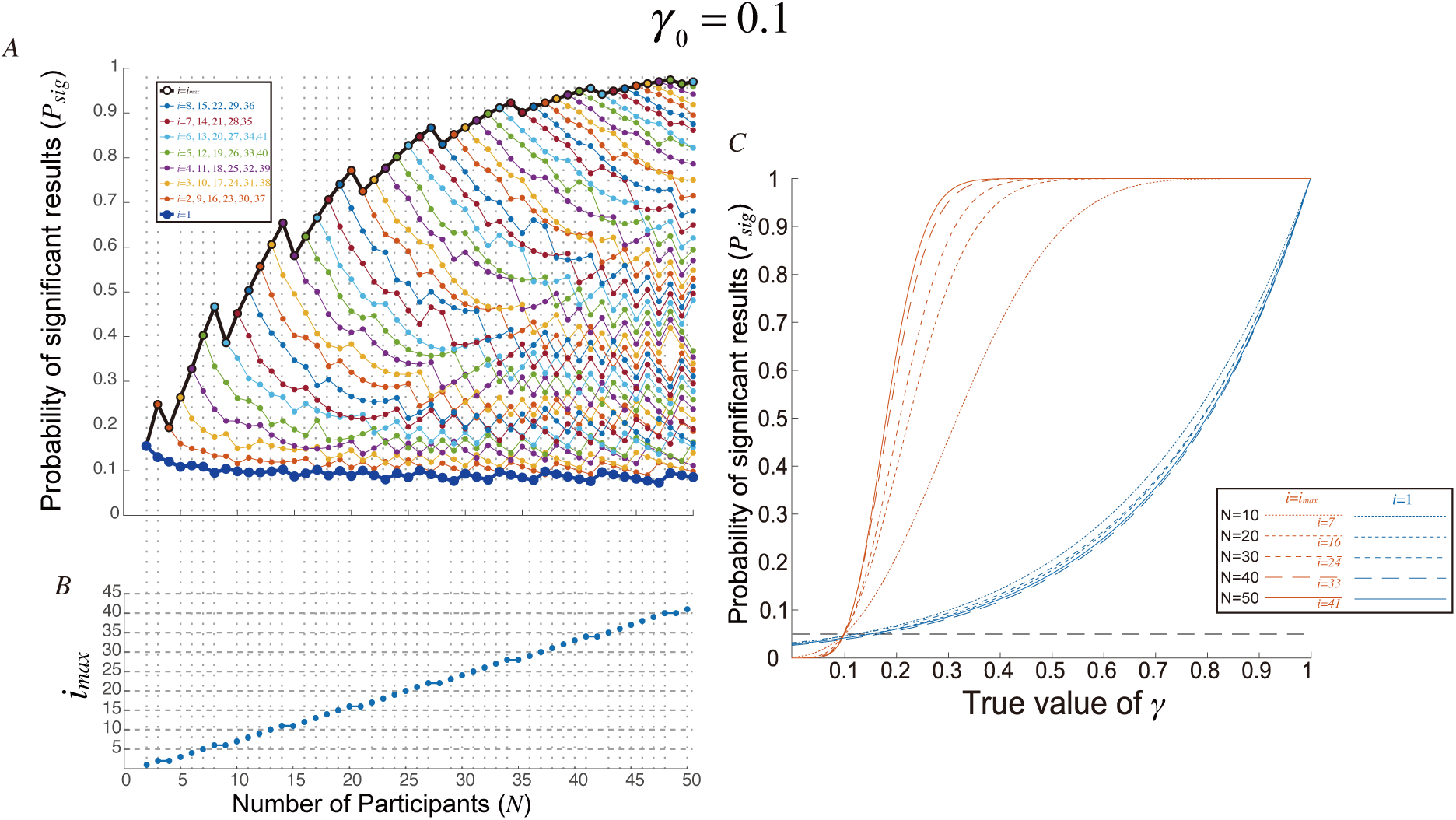
Results from the numerical calculation with *γ*_0_ = 0.1 is shown following the identical format to Figure 2. True gamma is set to 0.3. The other parameters are identical to those reported in the main text, i.e. *N*_*trial*_ = 1,000, *P*_*correct*+_ = 0.9, *α* = 0.05.

**Supplementary Figure S2:**
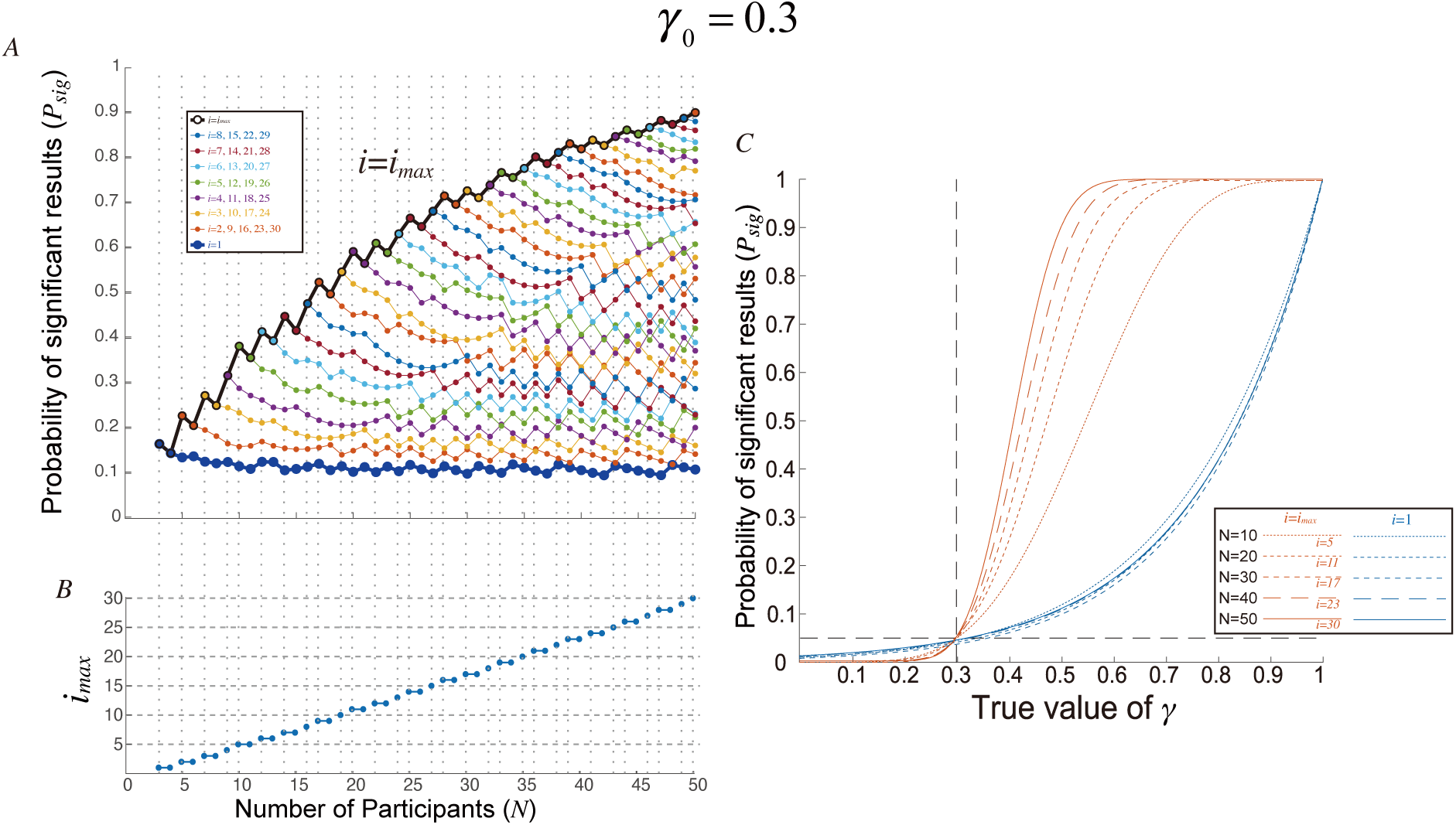
Results from the numerical calculation with *γ*_0_ = 0.0 is shown following the identical format to Figure 2. True gamma is set to 0.5. The other parameters are identical to those reported in the main text, i.e. *N*_*trial*_ = 1,000, *P*_*correct*+_ = 0.9, *α* = 0.05.

**Supplementary Figure S3:**
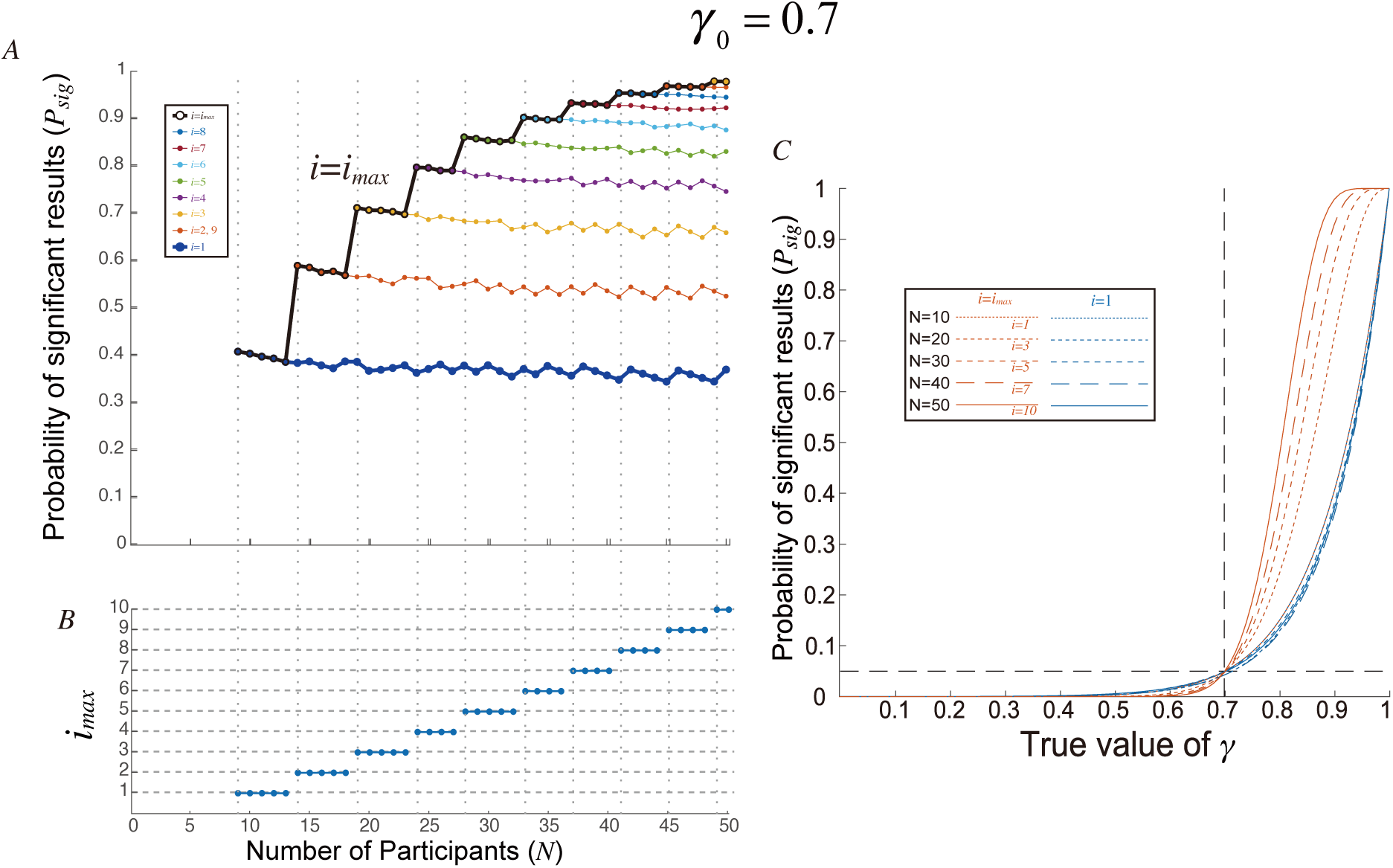
Results from the numerical calculation with *γ*_0_ = 0.7 is shown following the identical format to Figure 2. True gamma is set to 0.9. The other parameters are identical to those reported in the main text, i.e. *N*_*trial*_ = 1,000, *P*_*correct*+_ = 0.9, *α* = 0.05.

**Supplementary Figure S4:**
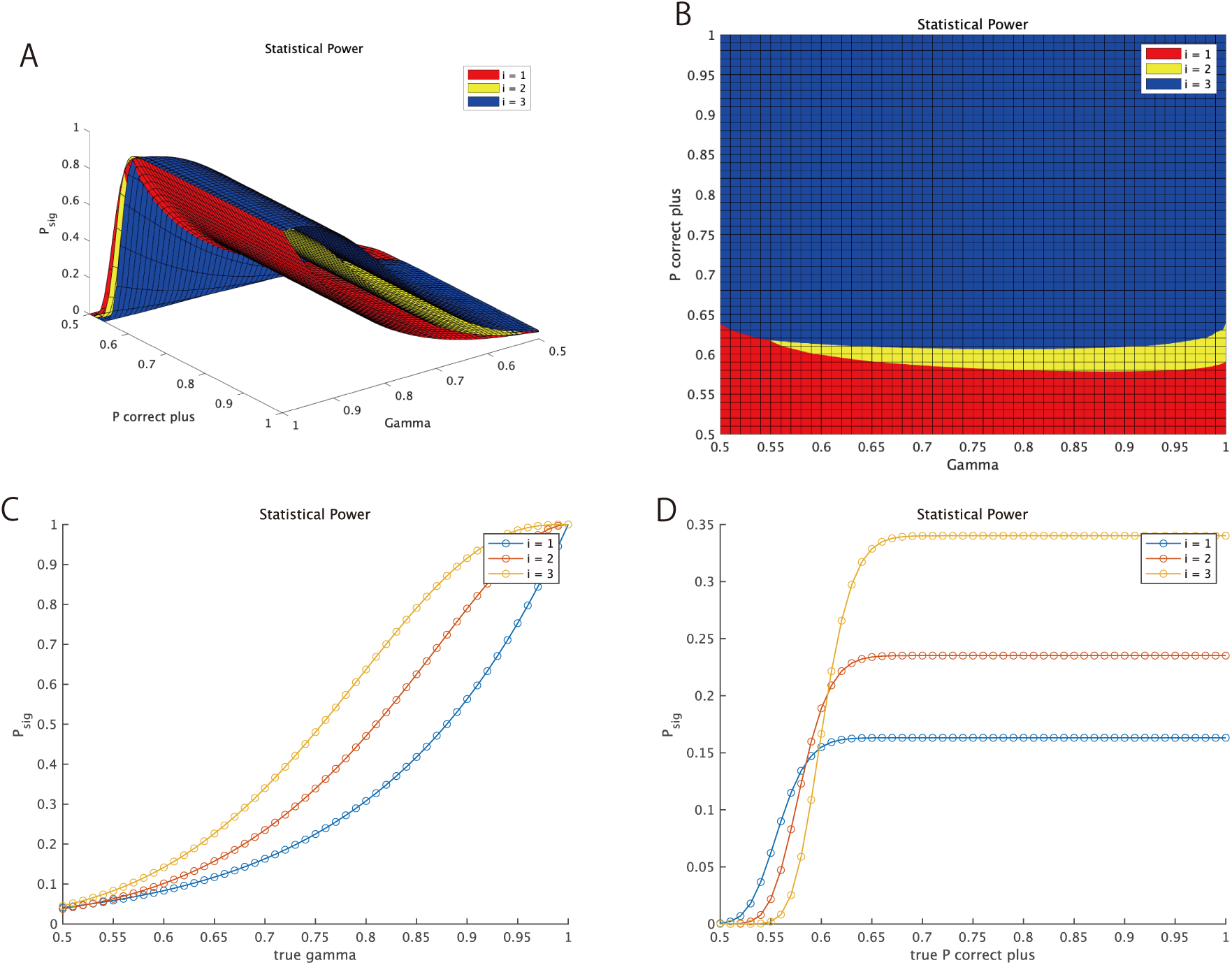
Statistical power expectation for the decision of *i* in the empirical experiment (*N* = 12, *N*_*trial*_ = 150). I set the predetermined threshold parameters as *γ*_0_ = 0.5 and *α* = 0.05. A: Expected statistical power (*P*_*sig*_; z-axis) is plotted against *P*_*correct*+_ (x-axis, from 0.5 to 1 with 0.01 steps) and *γ* (y-axis, from 0.5 to 1 with 0.01 steps) for *i* = 1, *i* = *2* and *i* = 0 (*i*_*max*_). B: Right from above view. Red, yellow and blue regions indicate that *i* = 1, 2 and 0 provides maximum statistical power. C: *P*_*correct*+_ is fixed at 0.7 and expected statistical power (*P*_*sig*_; y-axis) is plotted against *γ* (x-axis). D: *γ* is fixed at 0.7 and expected statistical power (*P*_*sig*_; y-axis) is plotted against *P*_*correct*+_ (x-axis). All panels are illustrated by the MATLAB program “`decide_i.m`.”

**Supplementary Figure S5:**
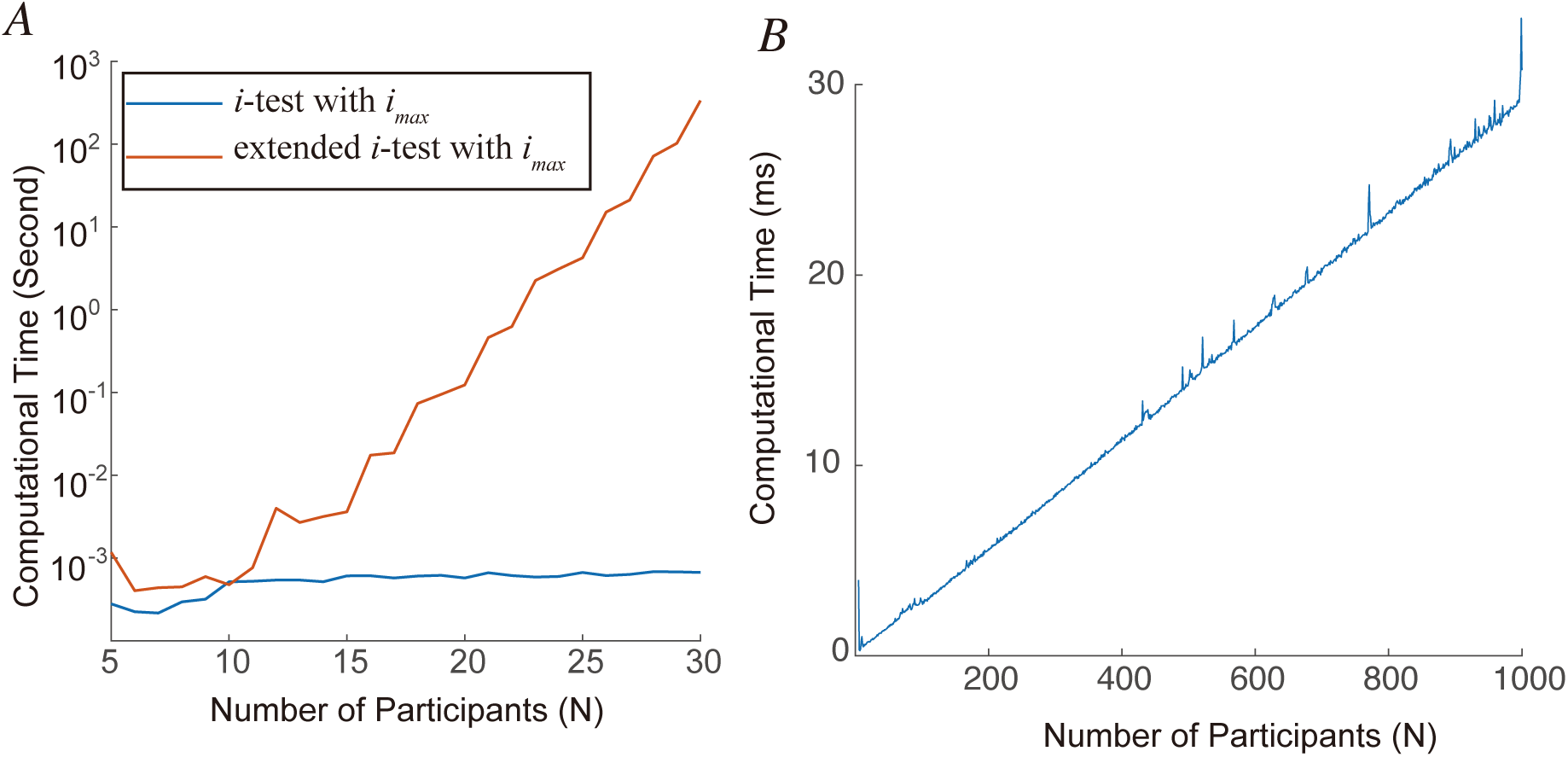
Benchmark test results on MATLAB 2018a, 64 bit, Linux, with 4 core of 3.5GHz CPU (Intel Xeon). **A**: Computational time for original i-test with maximum *i* (blue) and the extended i-test without identical distribution assumption (Appendix A; red) is plotted against the number of participants *N*. The vertical axis is log-scaled. **B**: Computational time for i-test with maximum *i*. The vertical axis is linear scaled.

**Supplementary Figure S6:**
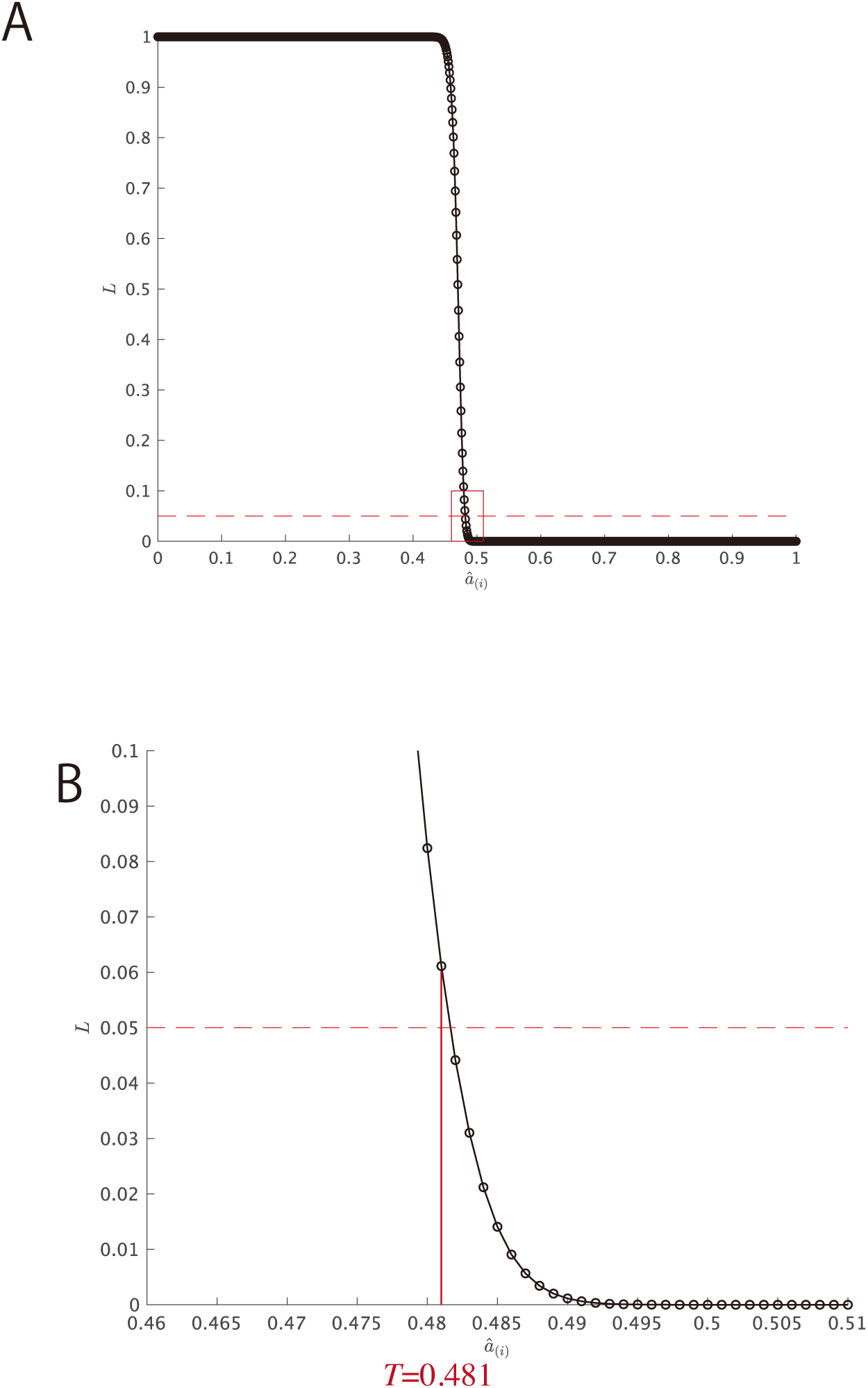
Calculation of the threshold *T*. The parameters are as follows; *α* = 0.05, *γ*_0_ = 0.5, *N* = 50, *i*=1, *N*_*trial*_ = 1,000, *P*_*correct*+_ = 0.9 and *γ* = 0.8. A: *L* is plotted against each value of *a*_(*i*)_. Note that *a*_(*i*)_ can take discrete values; from 0 to 1 with 0.001 steps. Each black circle corresponds to each value of *a*_(*i*)_. Horizontal dotted line indicates the statistical threshold *α* = 0.05. B: Zoom-up view of the region indicated by the red square in Panel A. Horizontal dotted line indicates the statistical threshold *α* = 0.05. Results indicate that *L* is smaller than *α* when and only when *a*_(*i*)_ is larger than 0.481 (*T* = 0.481).

## Reference

Faul, F., Erdfelder, E., Buchner, A., & Lang, A. G. 2009. Statistical power analyses using G* Power 3.1: Tests for correlation and regression analyses. Behavior research methods, 41(4), 1149–1160.

## References

Allefeld, C., Görgen, K., Haynes, J.D., 2016. Valid population inference for information-based imaging: From the second-level t-test to prevalence inference. Neuroimage. 141, 378–392. DOI: 10.1016/j.neuroimage.2016.07.040

Allefeld, C., Haynes, J.D. 2014. Searchlight-based multi-voxel pattern analysis of fMRI by cross-validated MANOVA. Neuroimage. 89, 345–357. DOI: 10.1016/j.neuroimage.2013.11.043

Brodersen, K.H., Ong, C.S., Stephan, K.E., Buhmann, J.M. The balanced accuracy and its posterior distribution. In Pattern Recognition (ICPR), 2010 20th international conference on (pp. 3121–3124). IEEE. DOI: 10.1109/ICPR.2010.764

Brodersen, K.H., Mathys, C., Chumbley, J.R., Daunizeau, J., Ong, C.S., Buhmann, J.M., Stephan, K.E., 2012. Bayesian mixed-effects inference on classification performance in hierarchical data sets. J. Mach. Learn. Res. 13, 3133–3176.

Deuker, L., Olligs, J., Fell, J., Kranz, T.A., Mormann, F., Montag, C., Reuter, M., Elger, C.E., Axmacher, N., 2013. Memory consolidation by replay of stimulus-specific neural activity. J. Neurosci. 33(49), 19373–19383. DOI: 10.1523/JNEUROSCI.0414-13.2013

Etzel, J.A., Zacks, J.M., Braver, T.S. 2013. Searchlight analysis: promise, pitfalls, and potential. Neuroimage. 78, 261–269. DOI: 10.1016/j.neuroimage.2016.07.040

Friston, K.J., Holmes, A.P., Poline, J.B., Grasby, P.J., Williams, S.C.R., Frackowiak, R.S., Turner, R., 1995. Analysis of fMRI time-series revisited. Neuroimage. 2(1), 45–53. DOI: 10.1006/nimg.1995.1023

Friston, K.J., Holmes, A.P., Price, C.J., Büchel, C., Worsley, K.J., 1999. Multisubject fMRI studies and conjunction analyses. Neuroimage. 10(4), 385–396. DOI: 10.1006/nimg.1999.0484

Fuentemilla, L., Penny, W.D., Cashdollar, N., Bunzeck, N., Düzel, E., 2010. Theta-coupled periodic replay in working memory. Curr. Biol. 20(7), 606–612. DOI: 10.1016/j.cub.2010.01.057

Gallivan, J.P., McLean, D.A., Valyear, K.F., Pettypiece, C.E., Culham, J.C., 2011. Decoding action intentions from preparatory brain activity in human parieto-frontal networks. J. Neurosci. 31, 9599–9610. DOI: 10.1523/JNEUROSCI.0080-11.2011

Gallivan, J.P., McLean, D.A., Flanagan, J.R., Culham, J.C., 2013. Where one hand meets the other: limb-specific and action-dependent movement plans decoded from preparatory signals in single human frontoparietal brain areas. J. Neurosci. 33, 1991–2008. DOI: 10.1523/JNEUROSCI.0541-12.2013

Gilbert, S.J., Fung, H., 2018. Decoding intentions of self and others from fMRI activity patterns. Neuroimage. 172, 278–290. DOI: 10.1016/j.neuroimage.2017.12.090

Good, P., 2000. Permutation tests: A practical guide to resampling methods for testing hypotheses. Springer

Haxby, J.V., Connolly, A.C., Guntupalli, J.S., 2014. Decoding neural representational spaces using multivariate pattern analysis. Annu. Rev. Neurosci. 37, 435–456. DOI: 10.1146/annurev-neuro-062012-170325

Hirose, S., Nambu, I., Naito, E., 2015. An empirical solution for over-pruning with a novel ensemble-learning method for fMRI decoding. J. Neurosci. Methods. 239, 238–245. DOI: 0.1016/j.jneumeth.2014.10.023

Hirose, S., Nambu, I., Naito, E., 2018. Cortical activation associated with motor preparation can be used to predict the freely chosen effector of an upcoming movement and reflects response time: An fMRI decoding study. Neuroimage. 183, 584–596. DOI: 10.1016/j.neuroimage.2018.08.060

Jamalabadi, H., Alizadeh, S., Schönauer, M., Leibold, C., Gais, S., 2016. Classification based hypothesis testing in neuroscience: Below*-*chance level classification rates and overlooked statistical properties of linear parametric classifiers. Hum. Brain. Mapp. 37(5), 1842–1855. DOI: 10.1002/hbm.23140

Kriegeskorte, N., Goebel, R., Bandettini, P. 2006. Information-based functional brain mapping. Proc. Natl. Acad. Sci. 103(10), 3863–3868. DOI: 10.1073/pnas.0600244103

Kowalczyk, A. Classification of anti-learnable biological and synthetic data. In European Conference on Principles of Data Mining and Knowledge Discovery (pp. 176–187). Springer, Berlin, Heidelberg. DOI: 10.1007/978-3-540-74976-9_19

Kowalczyk, A., Greenawalt, D.M., Bedo, J., Duong, C., Raskutti, G., Thomas, R.J., Phillips, W.A., 2007. Large Validation of Anti-learnable Signature in Classification of Response to Chemoradiotherapy in Esophageal Adenocarcinoma Patients. The First International Symposium on Optimization and Systems Biology (OSB’07).

Nambu, I., Hagura, N., Hirose, S., Wada, Y., Kawato, M., Naito, E., 2015. Decoding sequential finger movements from preparatory activity in higher*-*order motor regions: a functional magnetic resonance imaging multi*-*voxel pattern analysis. Eur. J. Neurosci. 42, 2851–2859. DOI: 10.1111/ejn.13063

Nili, H., Wingfield, C., Walther, A., Su, L., Marslen-Wilson, W., Kriegeskorte, N. 2014. A toolbox for representational similarity analysis. PLoS Comput Biol. 10(4), e1003553. DOI: 10.1371/journal.pcbi.1003553

Nishida, S., Nishimoto, S., 2018. Decoding naturalistic experiences from human brain activity via distributed representations of words. Neuroimage. 180, 232–242. DOI: 10.1016/j.neuroimage.2017.08.017

Noirhomme, Q., Lesenfants, D., Gomez, F., Soddu, A., Schrouff, J., Garraux, G., Luxen, A., Phillips, C., Laureys, S., 2014. Biased binomial assessment of cross-validated estimation of classification accuracies illustrated in diagnosis predictions. Neuroimage Clin. 4, 687–694. DOI: 10.1016/j.nicl.2014.04.004

Ojala, M., Garriga, G.C., 2010. Permutation tests for studying classifier performance. J. Mach. Learn. Res. 11(Jun), 1833–1863.

Roadknight, C., Aickelin, U., Qiu, G., Scholefield, J., Durrant, L. Supervised learning and anti-learning of colorectal cancer classes and survival rates from cellular biology parameters. In Systems, Man, and Cybernetics (SMC), 2012 IEEE International Conference. DOI: 10.1109/ICSMC.2012.6377825

Rosenblatt, J.D., Vink, M., Benjamini, Y., 2014. Revisiting multi-subject random effects in fMRI: Advocating prevalence estimation. Neuroimage. 84, 113–121. DOI: 10.1016/j.neuroimage.2013.08.025

Spiridon, M., Kanwisher, N., 2002. How distributed is visual category information in human occipito-temporal cortex? An fMRI study. Neuron. 35(6), 1157–1165. DOI: 10.1016/S0896-6273(02)00877-2

Stelzer, J., Chen, Y., Turner, R., 2013. Statistical inference and multiple testing correction in classification-based multi-voxel pattern analysis (MVPA): random permutations and cluster size control. Neuroimage. 65, 69–82. DOI: 10.1016/j.neuroimage.2012.09.063

Stephan, K.E., Penny, W.D., Daunizeau, J., Moran, R.J., Friston, K.J., 2009. Bayesian model selection for group studies. Neuroimage. 46 (4), 1004–1017. DOI: 10.1016/j.neuroimage.2009.03.025

